# ENQUIRE RECONSTRUCTS AND EXPANDS CONTEXT-SPECIFIC CO-OCCURRENCE NETWORKS FROM BIOMEDICAL LITERATURE

**DOI:** 10.1101/2023.09.10.556351

**Authors:** Luca Musella, Xin Lai, Max Widmann, Julio Vera

**Affiliations:** Laboratory of Systems Tumor Immunology, Friedrich-Alexander-Universität Erlangen-Nürnberg, Deutsches Zentrum Immuntherapie, BZKF, and Uniklinikum Erlangen, Erlangen, Germany; Systems and Network Medicine Lab, Biomedicine Unit, Faculty of Medicine and Health Technology, Tampere University, Tampere, Finland

**Author notes:** To whom correspondence should be addressed.; Correspondence may also be addressed to Julio Vera.

## Abstract

The accelerating growth of scientific literature overwhelms our capacity to manually distil complex phenomena like molecular networks linked to diseases. Moreover, biases in biomedical research and database annotation limit our interpretation of facts and generation of hypotheses. ENQUIRE (Expanding Networks by Querying Unexpectedly Inter-Related Entities) offers a time- and resource-efficient alternative to manual literature curation and database mining. ENQUIRE reconstructs and expands co-occurrence networks of genes and biomedical ontologies from user-selected input corpora and network-inferred PubMed queries. The integration of text mining, automatic querying, and network-based statistics mitigating literature biases makes ENQUIRE unique in its broad-scope applications. For example, ENQUIRE can generate co-occurrence gene networks that reflect high-confidence, functional networks. When tested on case studies spanning cancer, cell differentiation and immunity, ENQUIRE identified interlinked genes and enriched pathways unique to each topic, thereby preserving their underlying diversity. ENQUIRE supports biomedical researchers by easing literature annotation, boosting hypothesis formulation, and facilitating the identification of molecular targets for subsequent experimentation.

**GRAPHICAL ABSTRACT:** 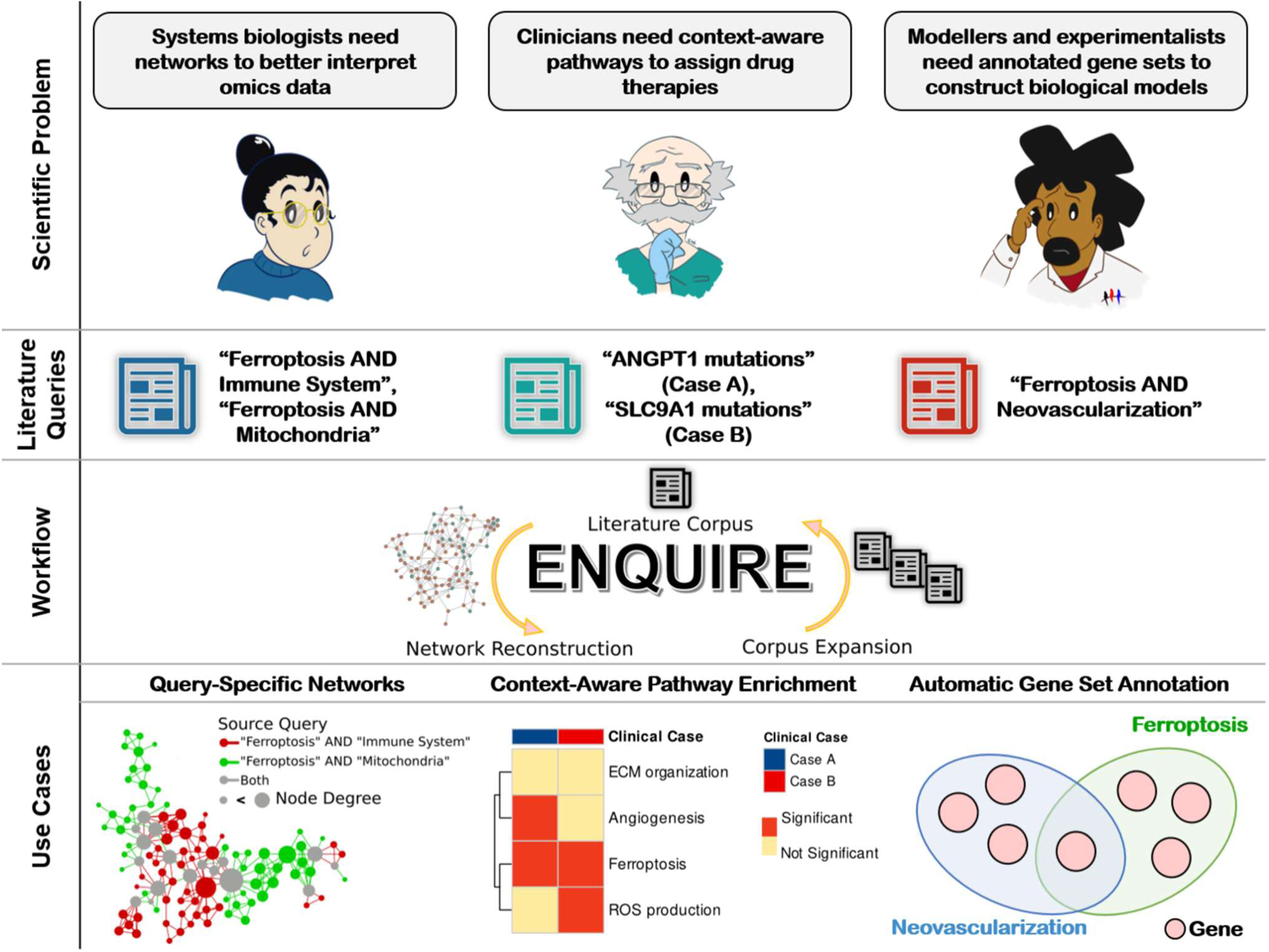

## INTRODUCTION

Curated gene networks are of high interest to prime the analysis of biomedical omics data, identification of disease-specific regulatory modules, and therapy-oriented studies like drug repurposing^1–4^. However, the growing biomedical literature corpus makes curation of biomolecular pathways challenging. Annotating molecular interactions from literature requires domain expertise, yet that same background knowledge could entail predispositions towards partial pictures of faceted biomedical problems^5^. In contrast, relation extraction from databases often omits the contextual information of gene interactions and can bias the results towards ubiquitously expressed, commonly investigated, and richly annotated genes^6–8^. This can make systematic comparisons of biomedical research topics inconclusive or unattractive from an expenditure perspective. Recently, there have been significant investments in the automatic annotation of scientific corpora. The knowledgebase immuneXpresso indexes textmined interactions among immune cells and cytokines^9^, while SimText provides a framework to interactively explore the content of a user-provided corpus of literature^10^. These and other tools rely on natural language processing methods like named-entity recognition^11^ (NER), part-of-speech recognition^12^, directionality assignment^13^, relationship detection, and co-occurrence scoring^14,15^. These efforts in biomedical text mining aim at detecting meta-features and co-occurrences in literature corpora. However, assessing the statistical significance and confidence level of a text-mined relation in dense, literature-based co-occurrence networks must be better addressed^16,17^. We find this striking, considering the well-documented reproducibility crisis^18–20^. In this context, we envisioned ENQUIRE (Expanding Networks by Querying Unexpectedly Inter-Related Entities) to achieve automatic reconstruction and expansion of biomedical co-occurrence networks from a user-defined PubMed literature corpus. ENQUIRE applies a state-of-the-art random graph model to retrieve context-specific, significant co-occurrences, i.e. dependent on the input corpus and its occurrence distribution of biomedical entities^21,22^. This distinctive element in our methodology allows ENQUIRE to control for literature biases. ENQUIRE processes scientific articles by extracting Medical Subject Headings (MeSH) and gene mentions from article abstracts, thus enriching gene-gene co-occurrence networks with gene-MeSH and MeSH-MeSH relations. ENQUIRE also automatically generates PubMed queries from connected biomedical entities in the network, contextually expanding the underlying corpus and, in turn, the co-occurrence network. To our knowledge, ENQUIRE is the first tool that integrates textmining, network reconstruction, and automatic literature querying into a single, resource efficient software. Here, we showcase ENQUIRE’s broad-scope applications and effectiveness in identifying relevant biomedical relations in different contexts and case scenarios.

## RESULTS

### A Tool to Generate Co-Occurrence Networks from Literature

ENQUIRE (Expanding Networks by Querying Unexpectedly Inter-Related Entities) is an algorithm that reconstructs and expands co-occurrence networks of *Homo sapiens* genes and biomedical ontologies (MeSH), using a corpus of PubMed articles as input. The method iteratively annotates MeSH and gene mentions from abstracts, statistically assesses their importance, and generates network-informed PubMed queries, until it obtains a connected network of genes and MeSH terms (or meets another exit condition). ENQUIRE’s pipeline implements a loop consisting of serial modules with the following structure (**Fig. 1**):

a) The user supplies an input literature corpus in the form of at least three PubMed identifiers (PMIDs).
b) The algorithm indexes the MeSH terms associated to the PMIDs listed. Next, their abstracts are parsed, and gene normalization is performed using a lookup table of gene aliases and abstract-specific blocklists of ambiguous terms.
c) ENQUIRE annotates and weights co-occurrences between gene and MeSH entities, accounting for the expected number of co-occurrences across the literature corpus.
d) The method selects significant co-occurrences and generates an undirected, simple graph, basing the test statistic on a random graph null model of unbiased mining of the input corpus.
e) Next, nodes are weighted, and “information-dense” maximal cliques, i.e. clusters of high-weight nodes all connected to each other, are selected to reconstruct network communities from the corresponding nodes.
f) ENQUIRE identifies optimal sets of community-connecting graphlets via an approximate solution to the “travelling salesman problem” (TSP).
g) Finally, the algorithm uses the entity nodes corresponding to the identified community-connecting graphlets into PubMed queries to find additional, relevant articles. Should ENQUIRE find new articles, their PMIDs are joined with the previous ones and automatically provided to module a), starting a new iteration.

**Fig. 1.**
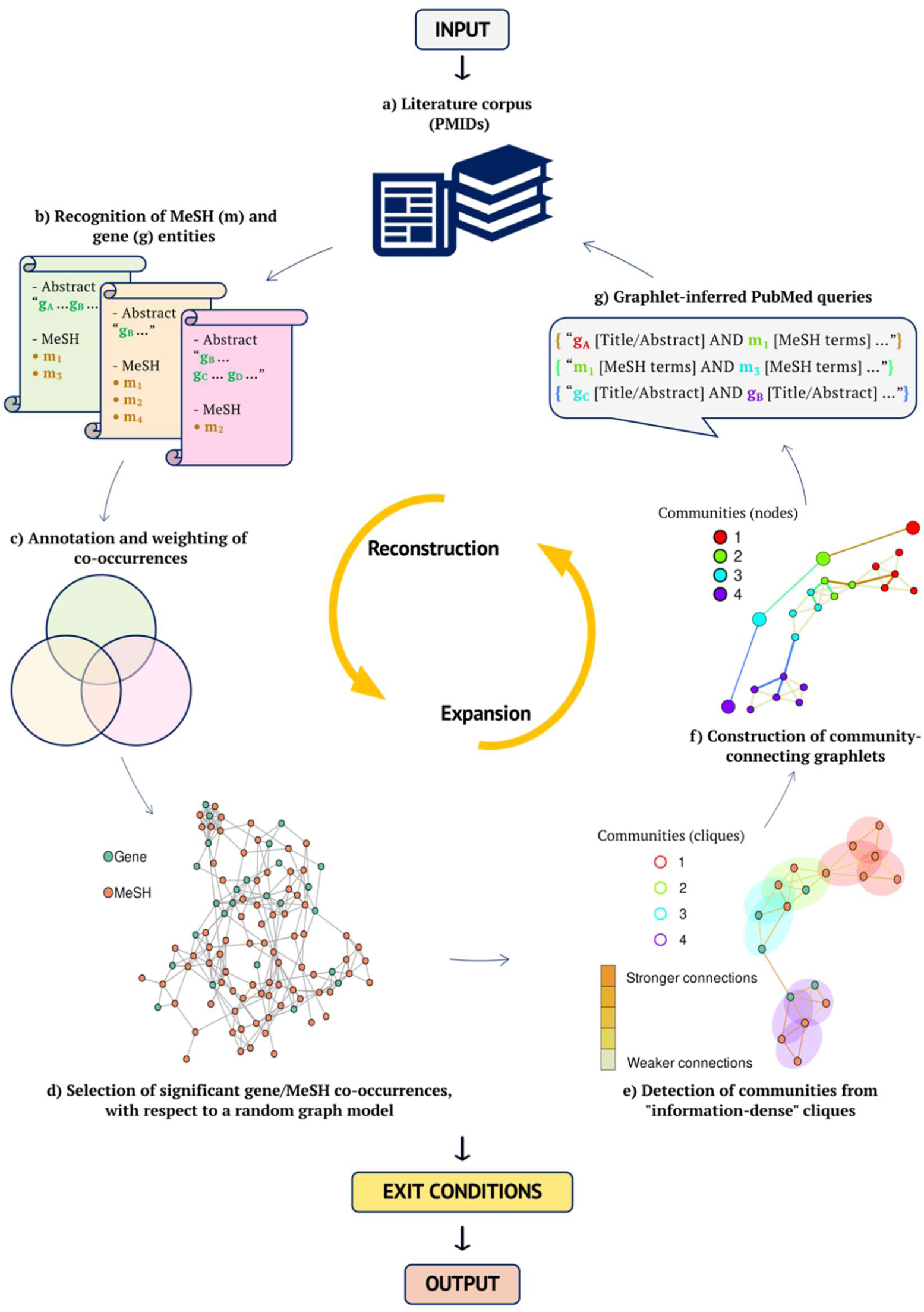
Overview of ENQUIRE methodology. ENQUIRE accepts a set of PubMed identifiers as input, together with optional, user-specified parameters. The pipeline iteratively orchestrates reconstruction and expansion of literature-derived co-occurrence networks, until an exit condition is fulfilled. Additional information about each alphabetically indexed module and output is provided in the **Mat.Met.** section. For a more detailed flowchart, see **Supp. Fig. 1**.

Whenever ENQUIRE reconstructs a network from the union of old and new PMIDs, the previously reconstructed network is joined with the new one. The joined network has recomputed edge and node weights in accordance to its expanded literature corpus and connectivity. The rationale is to prioritize the original reconstruction, while also leveraging the expanded literature corpus. Users can tune five options to tailor the workflow, namely: 1) Restricting the target entities to annotate genes or MeSH only – default: both; 2) representativeness threshold 𝑡 to disregard subgraphs characterized by poor overlap with the literature corpus – default: 1% overlap; 3) query size 𝑘 to control the number of entities that must be simultaneously used in a PubMed query – default: 4 entities; 4) query attempts 𝐴 to choose the number of attempts at connecting network communities by querying – default: 2 attempts; and 5) connectivity criterion 𝐾 to exclude newly found entities not having edges with nodes from 𝐾 communities previously generated at step (e) – default: 2 communities. ENQUIRE’s goal is to generate a gene/MeSH network and its respective gene- and MeSH-only subgraphs that individually consist of a single, connected component. The loop terminates if i) the network is empty after module d); ii) no clique can be found in step e); iii) the clique network consists of only one community; iv) all generated queries return empty results. With default parameters, ENQUIRE outputs node and edge lists of a gene/MeSH co-occurrence network and the respective gene- and MeSH-only subgraphs at each iteration. The final ENQUIRE results include additional tabulated data, graphics, and links to collected resources for subsequent analyses and reproducibility. For instance, it is possible to extract subsets of the literature corpus that support a gene/MeSH relation of interest and access the articles via hyperlinks redirecting to PubMed.

See **Supp. Fig. 1** and **Mat.Met.** for a comprehensive description of the algorithm.

### An Exemplary ENQUIRE Run

To showcase ENQUIRE, we set up a small-scale case study in which we looked for literature-based relationships between the immune system and ferroptosis, a form of programmed cell death^23^. We selected 27 papers obtained from the PubMed query *(“Ferroptosis”[MeSH terms] AND “Immune System”[MeSH terms]) NOT “review”[Publication Type]”* – queried on 14.04.23. We increased the number of attempts 𝐴 to 3, as we expected few query-matching PMID. The expansion process is depicted in **Fig. 2A**, using the Cytoscape package DyNet^24,25^. The original reconstructed network consists of four connected components. The first expansion led to additional, significant co-occurrences and newly found entities that connected the four components into a single one. The algorithm stopped after obtaining a single, connected gene/MeSH network and not finding additional query-matching PMIDs. Using up to 6 CPU cores, ENQUIRE finished in 16 minutes using less than 0.4 GB of RAM (**Supp. Fig. 2**). Next, we applied context-specific gene set annotation on the original gene/MeSH co-occurrence networks, as described in **Mat.Met.** We identified non-trivial, descriptive gene sets (**Fig. 2C**-left), including ferroptosis-dependent inflammation supported by immune-related adaptor proteins (blue, top left), antineoplastic effects of the ferroptosis-inducer sulfasalazine acting on the amino acid transport system (magenta), and cross-talk between ferroptosis and autophagy (pink), in accordance with previous findings^26–28^. We also performed context-aware pathway enrichment analysis using the gene-gene co-occurrence subgraphs and the approach described in **Mat.Met.** We summarized the results in **Fig. 2C**-right, which depicts 30 Reactome pathways whose adjusted p-values were below 5% FDR for at least one network, sorted by Reactome category. In the original network, we obtained enrichments of pathways centered around Toll-like receptor and MAP kinases signaling cascades (e.g. R-HSA-975138). In the expanded networks, the metabolic pathway *Glutathione conjugation* (R-HSA-156590) and additional innate immunity-related and programmed cell death pathways were enriched. Taken together, the ENQUIRE-generated output highlights potential molecular axes between iron-regulated cell death and proliferation, metabolism, and immune response^29–31^.

**Fig. 2.**
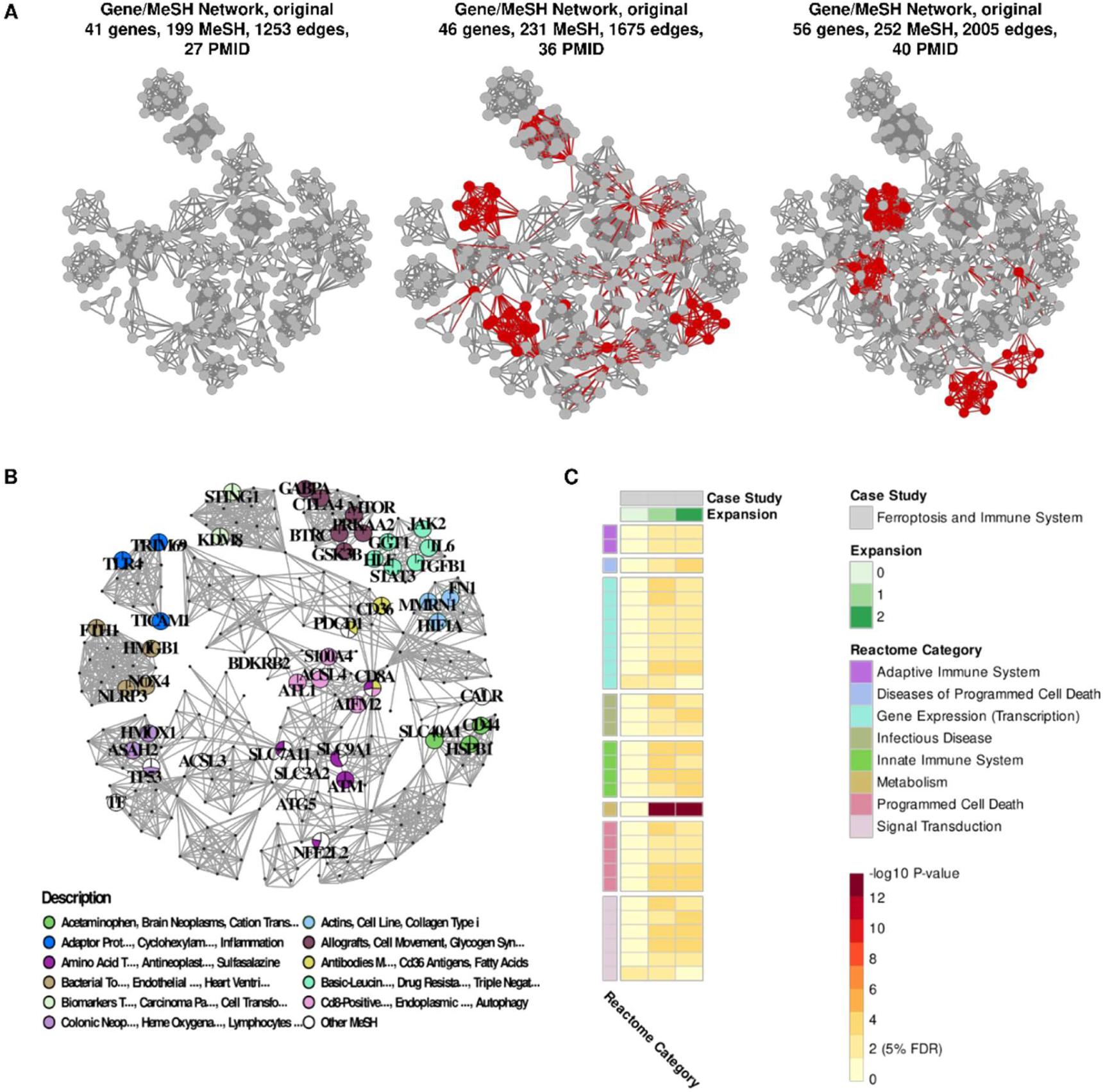
Example of ENQUIRE’s network reconstruction, expansion and post-hoc analyses. We used the PubMed identifiers (PMIDs) obtained from the query *(“Ferroptosis”[MeSH terms] AND “Immune System”[MeSH terms]) NOT “review”[Publication Type]* as input. **A**: visualization of ENQUIRE’s network expansion process. Newly found nodes and edges are indicated in red at each expansion. **B**: output of the automatic gene set reconstruction, using the original Gene/MeSH network as input and fuzzy c-means. For simplicity, only nodes referring to genes are enlarged and labelled, and a shortened description of computed gene sets of size 2 or bigger is provided. Sector sizes of the pie-chart-shaped nodes reflect their relative membership degree with respect to each cluster. **C**: topology-based enrichment analysis of Reactome pathways, using original and expanded networks, as described in the Methods section. 30 pathways whose adjusted p-value was significant in at least two networks are depicted. Reactome pathways are grouped based on “Top-Level Pathway” and “Disease” categories. FDR: Holm’s family wise error rate.

### ENQUIRE’s Gene Normalization Strategy is Precise and Efficient

ENQUIRE is intended to consume abstracts from studies in *H. sapiens* and *M. musculus*. We therefore evaluated ENQUIRE’s precision and recall using the abstracts in the NLM-Gene corpus mentioning at least one *M. musculus* or *H. sapiens* gene – 479 out of 550 entries^32^. ENQUIRE’s maximum F1 score is 0.747, corresponding to 0.822 precision and 0.683 recall, using as little as 0.36 GB of RAM and with speeds up to 0.03 seconds per abstract (**Table 1**). The Schwartz-Hearst abbreviation-definition detection algorithm improves precision of tokenization and normalization by 2%, without major loss in recall nor higher computational requirements^33^. In some use cases, it could be necessary to exclude gene mentions associated to cell entities, such as “CD8+ lymphocytes”. The scispaCy’s *en_ner_jnlpba_md* model removes unwanted gene-matching cell mentions, at the cost of about 2% reduction in recall^34^. It should be noted, however, that the latter metric is affected by the fact that gene mentions included in cell entities are counted as true positives in the NLM-Gene corpus. We also compared ENQUIRE’s performance to GNorm2, a state-of-the-art deep-learning model for gene entity recognition and normalization^35^. We tested ENQUIRE’s most resource-intensive configuration (both *en_ner_jnlpba_md* and *Schwartz-Hearst* modules enabled) against GNorm2’s implementation of Bioformer, a deep-learning model based on BERT, but 60% smaller in size^36^. **Table 2** shows that GNorm2 is considerably slower and has a higher resource usage than ENQUIRE. If ENQUIRE were to implement GNorm2 for gene normalization, this would impair its usage in scenarios with limited resources and computing time: for example, we verified that GNorm2 cannot be run on the CPU-based computer with 16GB of RAM used for the exemplary case study (**Supp. Fig. 3** and **Supp. Information**). In this terms, ENQUIRE’s *in-house* gene normalization is more suitable for textmining large input corpora on a variety of devices beyond CPU-based computer clusters.

**Fig. 3.**
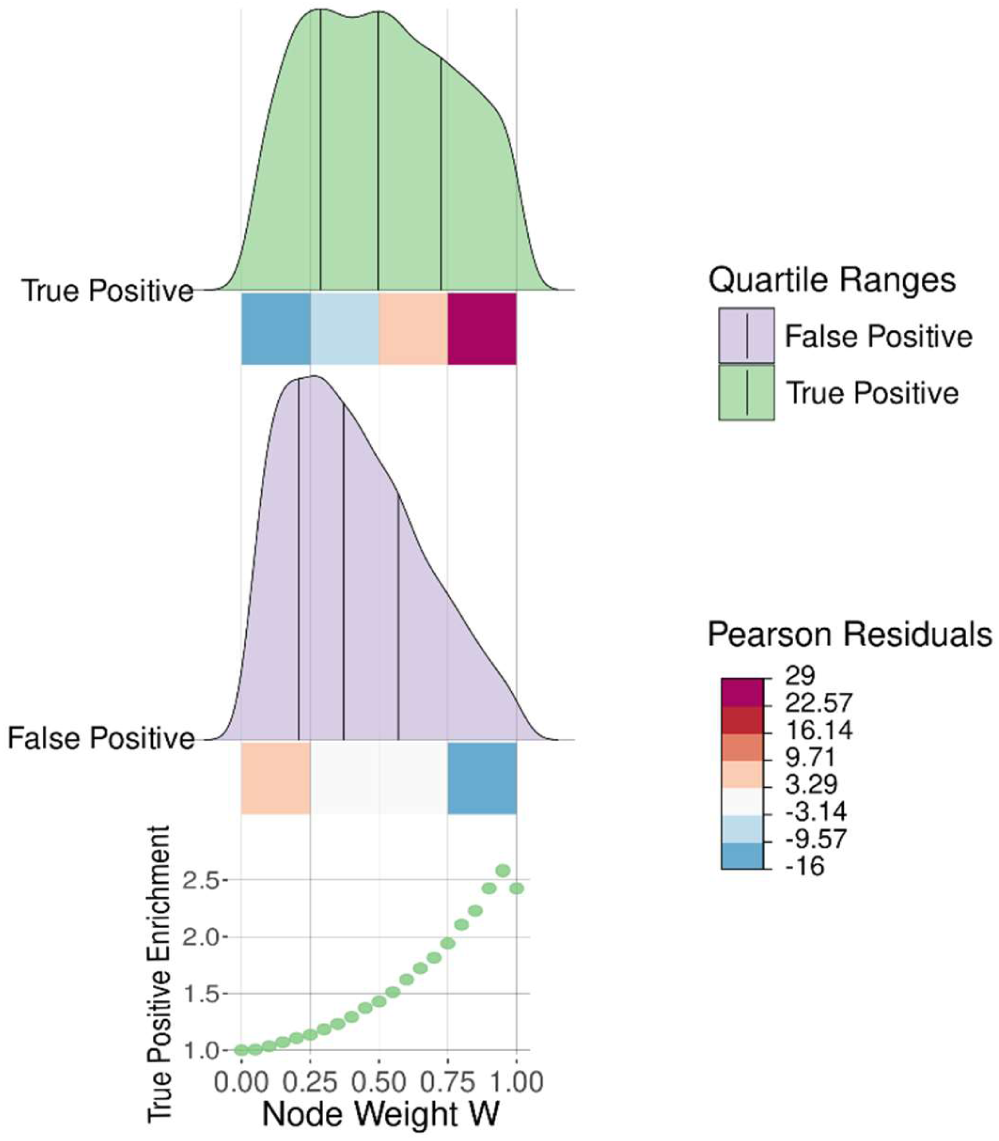
Node weight distribution of ENQUIRE-derived gene networks correlate with relevance to the input literature corpus. We defined true and false positives genes according to their presence or absence in a Reactome pathway, whose reference literature was used to retrieve gene mentions via ENQUIRE’s gene normalization and network reconstruction. The statistics shows the aggregated results from 720 Reactome-derived input corpora. The aggregated distributions for true and false positive genes are segmented into quartiles. We defined four ranges of the node score 𝑊, indicated by squares, whose colors reflect Pearson standardized residuals resulting from a significant chi-square statistic. The lower chart depicts the enrichment of true positive genes, after pruning ENQUIRE-derived networks based on different values of 𝑊. Values are relative to the original proportion of true positives.

**Table 1.** Selection of case studies for assessment of context resolution at the molecular pathway level. We obtained PubMed queries by “AND” concatenation of up to three MeSH terms and further filters to retrieve review articles only. The Corpus sizes refer to the non-redundant union of publications cited by three independent review articles, reported under the “References” column.

| Major Topic | Case Study<br>(abbreviation) | PubMed Query |  |  | Corpus<br>size | References<br>(PMID) |
| --- | --- | --- | --- | --- | --- | --- |
|  |  | MeSH 1 | MeSH 2 | MeSH 3 |  |  |
| Signal transduction in solid tumors | Melanoma (MM-ST) | Signal transduction | Melanoma |  | 944 | 25587943, 32605090, 34924562 |
|  | Uveal melanoma (UM-ST) | Signal transduction | Uveal neoplasms |  | 218 | 25296731, 25113308, 28223438 |
|  | Colorectal cancer (COL) | Signal transduction | Colorectal neoplasms |  | 556 | 34884633, 34742312, 35836256 |
|  | Breast cancer (BRE-ST) | Signal transduction | Breast neoplasms |  | 522 | 29455658, 31752925, 32245065 |
| Macrophage’s signal transduction in disease | Macrophage signal transduction upon infection (MP-ST) | Signal transduction | Macrophages | Mycobacterium tuberculosis | 470 | 32849525, 33558322, 34502407 |
|  | Tumor-associated Macrophages (MP-TA) | Signal transduction | Tumor associated macrophages |  | 386 | 33365025, 35844605, 35740975 |
| Antigen presentation in autoimmune diseases | Inflammatory bowel disease (IBD-AP) | Antigen presentation | Inflammatory bowel diseases |  | 445 | 28534191, 33584726, 33800865 |
|  | Rheumatoid arthritis (RA_AP) | Antigen presentation | Arthritis, rheumatoid |  | 452 | 27225300, 28451787, 30589082 |
|  | Psoriasis (PSO-AP) | Antigen presentation | Psoriasis |  | 435 | 26215033, 29316717, 33050592 |
| Oligodendrocyte differentiation | Oligodendrocyte (ODC) | Cell differentiation | Oligodendroglia |  | 355 | 24979526, 30770136, 31614602 |
| Positive control | All case studies (CTR) | All queries (“OR” concatenation) |  |  | 3606 | All of the above |

**Table 2.**
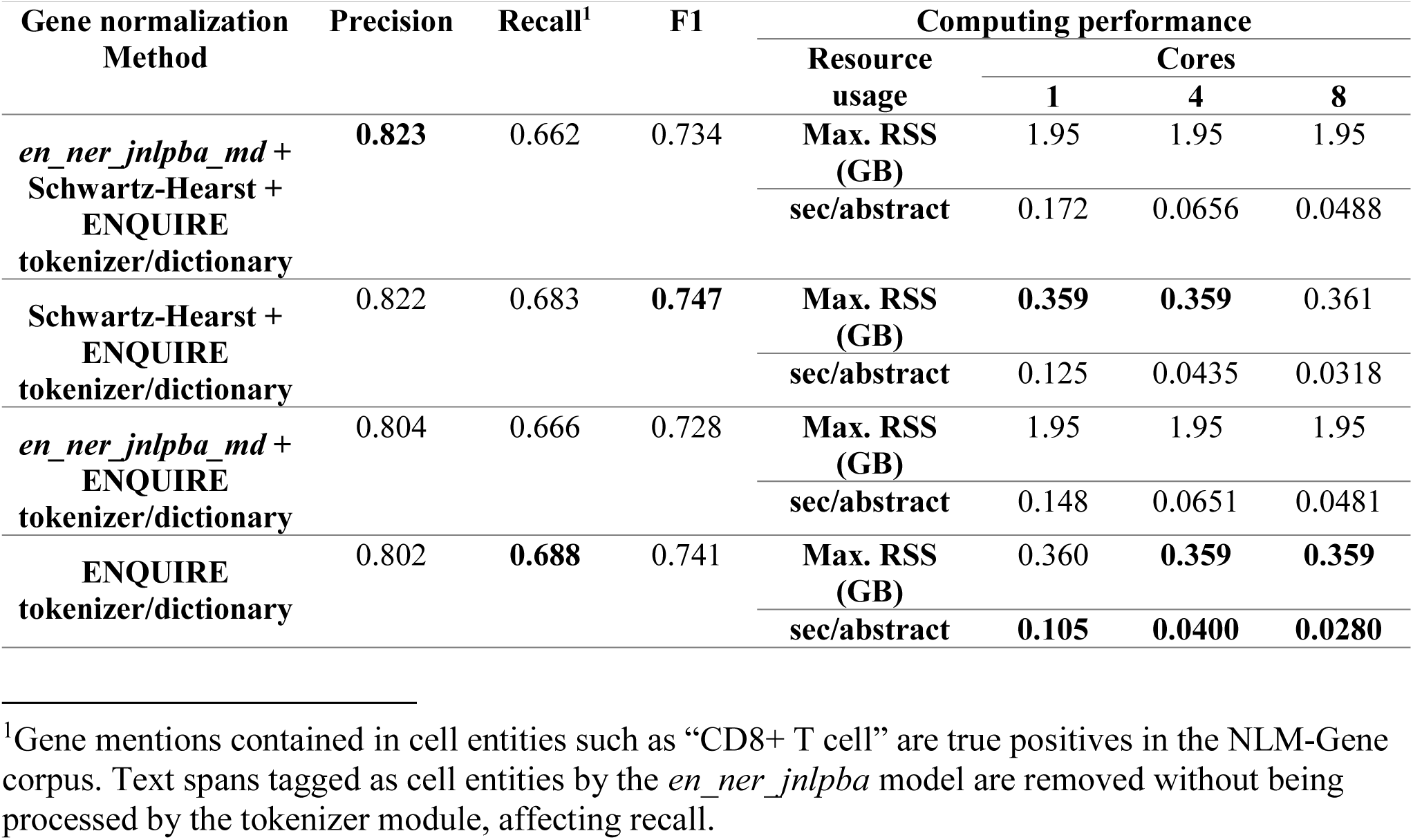
Performance of ENQUIRE’s gene normalization algorithm. The gene normalization task is here defined as detecting at least one gene alias per unique reference gene mentioned in an abstract. Precision, recall, and their harmonic mean (F1) are based on annotated abstracts from the NLM-Gene corpus containing at least one mention to a *H. sapiens* or *M. musculus* gene (479 abstracts). We ran the computations on a Linux computer with 20 CPUs (3.1 GHz) and 252 GB of RAM. Up to 8 cores were used for parallelization. We tested different gene normalization methods by adding or removing filters for excluding predicted cell entities (*en_ner_jnlpba_md*) and ambiguous abbreviation-definition pairs (Schwartz-Hearst). Maximum RAM usage is measured as resident set size (RSS). Estimated time in seconds per abstract (sec/abstract) also accounts for loading the gene alias lookup table and machine learning models. The best value for each parameter setting is highlighted in bold.

### ENQUIRE Networks Support Ranking of Genes Relevant to the Input Literature

To evaluate ENQUIRE’s ability in inferring genes relevant to the input corpus, we extracted *H. sapiens* pathways, their belonging genes, and corresponding primary literature references from the Reactome Graph Database^37^. We used the lists of references as inputs and performed a single gene entity-restricted co-occurrence network reconstruction for each pathway. Out of 967 examined pathways, ENQUIRE successfully reconstructed a gene co-occurrence network from the reference literature of 733 of them. We evaluated the effect of input corpus size, pathway size and average entity co-occurrence per paper on the accuracy of the resulting networks (**Table 3**). As expected, precision and recall show opposite Spearman’s correlation trends concerning corpus and pathway sizes, but average gene-gene co-occurrence per article appears uncorrelated. The negative correlation between corpus size and precision is -0.18, suggesting a low impact of large input corpora on the output. Next, we explored if the ENQUIRE-computed weight 𝑊, an aggregated measure of network centrality and literature support of its connections, is a useful measure of gene relevance regarding the input corpus (**Mat.Met.**). To this end, we analyzed the above-mentioned gene-scope co-occurrence networks. In **Fig. 3**, we compare the pan-pathway-aggregated distributions of true-positive (top panel) and false-positive (middle panel) ENQUIRE-derived genes as a function of 𝑊 (x-axis). We subdivided the distribution into four evenly spaced intervals, performed a chi-square test of independence, which resulted to be significant, and extracted the standardized Pearson residuals for true positives and false positives (colored boxes beneath the distributions). True positives tend to have higher node weights than false positives. An over-representation of node weights higher than 0.75 is observed in the true-positive distribution, as indicated by the color gradient in Pearson residuals. This suggests one can use the node weights 𝑊 to rank a set of ENQUIRE-derived genes based on their relevance to the literature corpus in question.

**Table 3.**
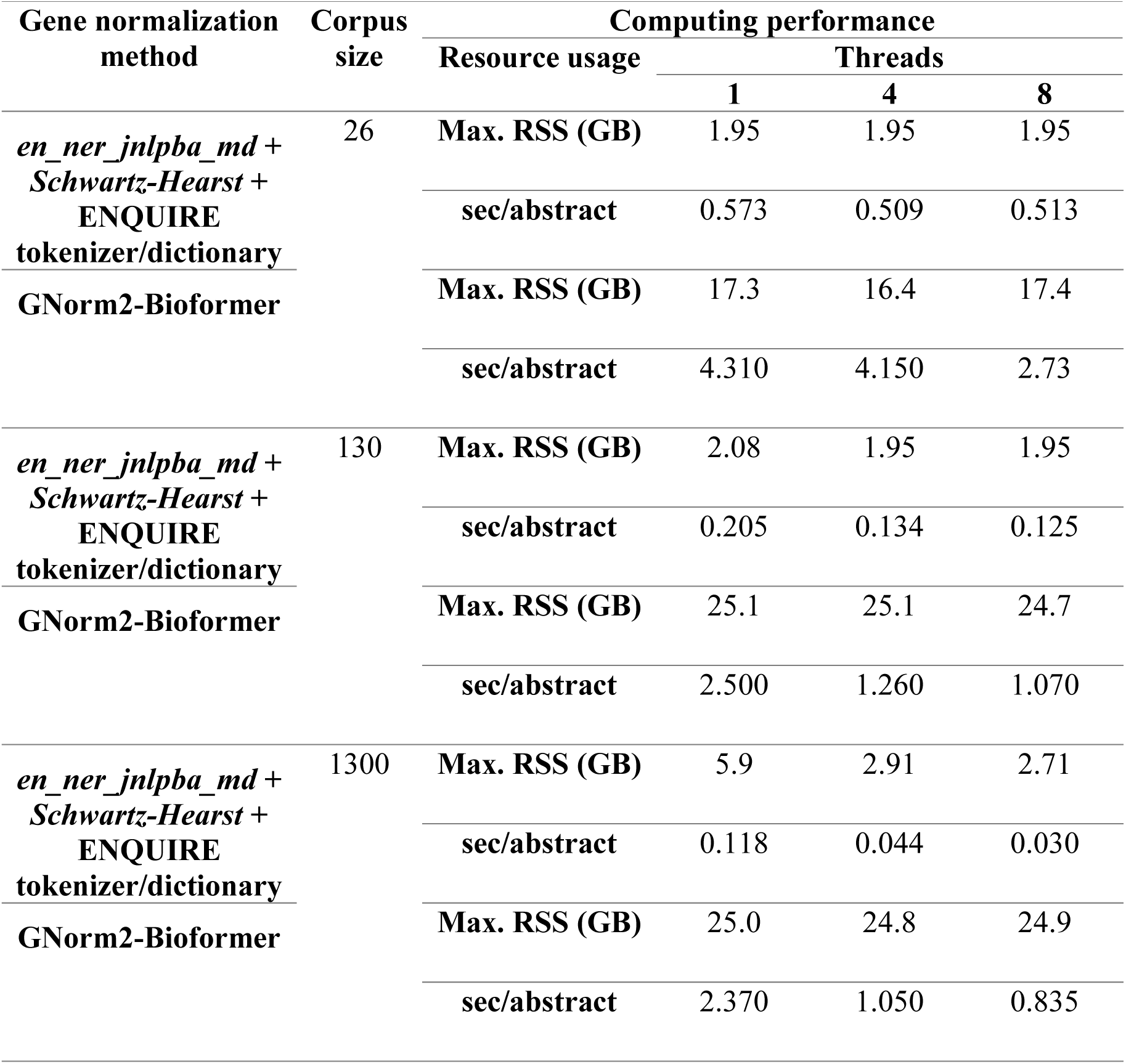
Differences in computing performance between ENQUIRE’s gene normalization algorithm and GNorm2-Bioformer. We ran the computations on a Linux computer with 20 CPUs (3.1 GHz) and 252 GB of RAM. Up to 8 cores were used for parallelization. Maximum RAM usage was measured as resident set size (RSS). Estimated time in seconds per process abstract (sec/abstract) also accounts for loading of gene alias lookup table and machine learning models.

### ENQUIRE Recovers Genes with High Chances of Showing Biochemical Interrelations

We hypothesized that ENQUIRE-derived gene co-occurrence networks could be enriched in molecular gene-gene interactions annotated in databases. To test this, we queried PubMed with all possible cross-pairs of *Diseases* and *Genetic Phenomena* MeSH terms. We further processed the 3098 queries that retrieved 50-500 matching PMIDs and extracted their gene-gene co-occurrence networks obtained after one network reconstruction. We then inspected whether their respective protein-coding genes can produce significant functional association networks based on STRING’s protein-protein interaction (PPI) database^38^ (see **Mat.Met.**). **Table 4** indicates that for 1336 (43.1%) MeSH pairs, both ENQUIRE and STRING generated a minimal network with at least three genes and two edges. In a subset of 733 network with degree sequences allowing at least ten different graph realizations, we assessed ENQUIRE’s capability of reflecting functional interactions. Then, we then generated two empirical random probability distributions for STRING’s edge count and DeltaCon similarity score^39^ (see **Mat.Met.**). Within the tested networks, 730 protein-coding gene networks (99.6%) produced a STRING network with a higher edge count than 95% of equal-sized random STRING networks (PPI score). At the same time, 439 networks (59.9%) showed concordance with STRING-derived PPI networks based on statistically significant DeltaCon similarities. After p-value adjustment, (1% FDR, **Table 3**), 722 (98.5%) and 344 (46.9%) ENQUIRE networks still show significantly high PPI scores and DeltaCon similarities, respectively. To evaluate the effect of network size, we subdivided the 733 suitable networks into quartiles based on their node number and mapped the respective unadjusted p-value distributions of the above-described test sets. The edge-count-associated p-values increased with network size (**Fig. 4A**). At the same time, the observed DeltaCon similarity values monotonically decrease with network size (**Table 5**). This is in accordance with DeltaCon’s implementation of edge importance and zero-property^39^, as differences in edge counts and number of connected components between ENQUIRE and STRING increase with the number of nodes. Nevertheless, we did not find a negative correlation between network size and p-values of observed DeltaCon similarities; instead, the quartile corresponding to the largest network also shows the largest relative proportion of significant adjusted p-values (**Fig. 4B**). Taken together, our results suggest that ENQUIRE generates networks that frequently contain established, high-confidence functional relations.

**Fig. 4.**
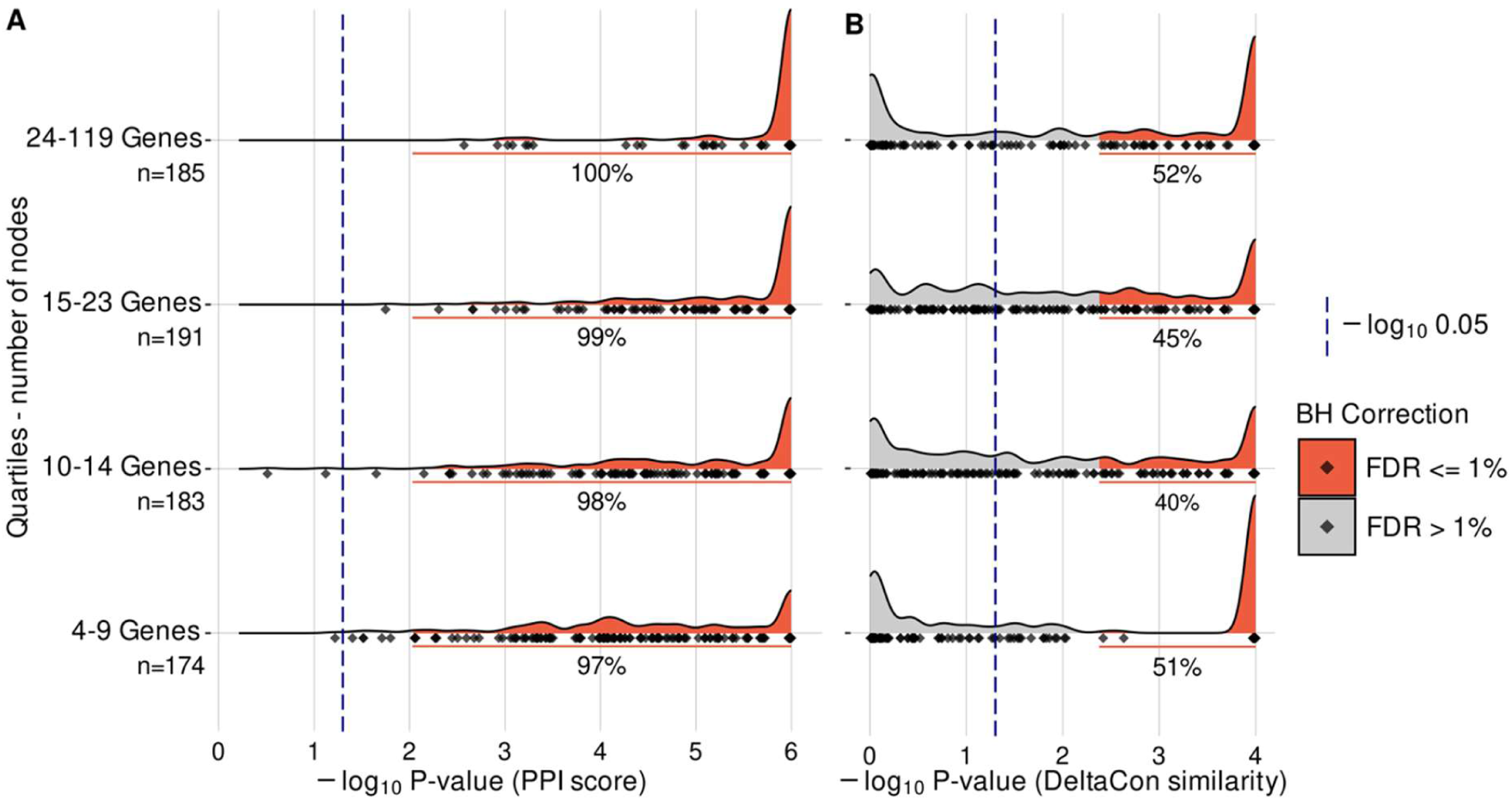
Protein-coding genes from ENQUIRE-generated graphs significantly share functional associations. Panels (A) and (B) respectively report the unadjusted p-value density distributions of STRING-informed edge counts and DeltaCon similarities, arranged by number of protein-coding genes (network size). We used the *H. sapiens* functional association network from STRING to evaluate ENQUIRE-derived networks of protein-coding genes. We tested 733 networks having 10 or more possible network realizations given the observed degree sequence. For each observed network size and degree sequence of ENQUIRE-generated gene networks, 1,000,000 and 10,000 samples were respectively generated to perform a test statistic on the observed edge counts and DeltaCon similarities. See **Mat.Met.** for additional information. The 733 tested networks are apportioned into quartiles based on network size, and for each the exact size is indicated (n). Within each network size interval, grey and red areas respectively highlight insignificant and significant p-values with respect to a globally-applied Benjamini-Hochberg correction (BH), and a percentage is indicated for those below 1% FDR. Diamonds indicate the observed data.

**Table 4.**
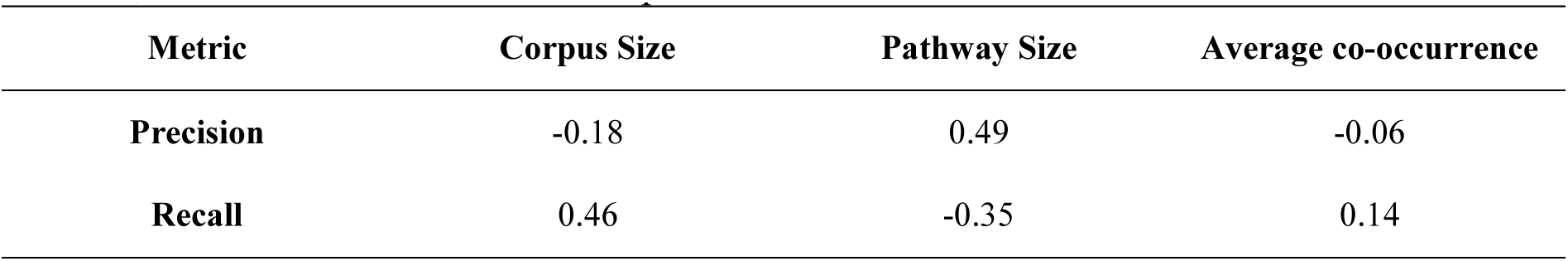
Effect of relevant covariates on quality indicators of ENQUIRE’s gene entity recognition. We evaluated the effect of corpus size (input), Reactome’s pathway size (number of genes to be retrieved) and average gene-gene co-occurrence per article, using Spearman’s correlation coe<u>fficients, for each measure.</u>

| Metric | Corpus Size | Pathway Size | Average co-occurrence |
| --- | --- | --- | --- |
| Precision | -0.18 | 0.49 | -0.06 |
| Recall | 0.46 | -0.35 | 0.14 |

**Table 5.** Relevant quality indicators of functional associations in 3098 case studies. PPI: protein-protein interaction score, as number of observed edges over the STRING-inferred network. FDR: false discovery rate, expressed in percentage. Percentages reported for PPI and DeltaCon significance independently refer to the set of 733 tested networks, i.e. those with 10 or more possible realizations with the same degree sequence as ENQUIRE-derived networks.

| Property | Subset |  | Raw count | Percentage over the preceding step | Percentage over total (3098) |
| --- | --- | --- | --- | --- | --- |
| Network topology | At least 3 genes and 2 edges in both ENQUIRE and STRING networks |  | 1336 | / | 43.1% |
|  | At least 10 possible realizations of the same degree sequence |  | 733 | 54.9% | 23.7% |
| Significance | Edge count p-value | < 0.05 | 730 | 99.6% | 23.6% |
|  |  | < 1% FDR | 722 | 98.5% | 23.3% |
|  | DeltaCon p-value | < 0.05 | 439 | 59.9% | 14.2% |
|  |  | < 1% FDR | 344 | 46.9% | 11.1% |

### ENQUIRE Improves the Context Resolution of Topology-Based Pathway Enrichment Analyses

We also analyzed ENQUIRE’s ability to generate and expand co-occurrence networks with distinctive biological and biomedical signatures by literature querying. In particular, we evaluated the context resolution of ENQUIRE-generated gene networks, i.e. their ability to preserve differences and similarities in gene mention content from different corpora. To this end, we applied the complete ENQUIRE pipeline with default parameters to a comprehensive set of case studies, spanning cancer, cell differentiation, innate immunity, autoimmune diseases, and a positive control (**Table 6**). Notice that each case study’s input corpus is a perfect subset of the positive control corpus, which corresponds to a Szymkiewicz-Simpson overlap coefficient (OC) of 100% - see **Mat.Met.**. Despite that, the positive control network does not always exhibit an OC of 100% with non-expanded networks, in terms of both nodes and edges (**Supp. Fig. 4**). This shows that ENQUIRE’s network reconstruction is sensitive to the input corpus. **Fig. 5A** depicts the expected dendrogram of the different case studies and respective expansions, based on their major topics and original input corpora. **Fig. 5B** shows the observed clustering based on ENQUIRE-informed, topology-based pathway enrichment analysis using KNet^40^ (see Post Hoc Analyses in **Mat.Met.** and **Supp. Fig. 2**). The 50 pathways with at least one significant, adjusted p-value (5% FDR) and highest p-value variances across case studies are depicted. The heat-map suggests that the case studies primarily cluster based on the affinities between their major topics, in agreement with the expected dendrogram. For example, pathways categorized under *Diseases of Metabolism*, *Diseases of Immune System*, and *Innate Immune System* are predominantly enriched in networks originated from the case study “Macrophage’s signal transduction during M. tuberculosis infection” (MP-ST) and the major topic “Antigen Presentation in Autoimmune Diseases”. Similarly, some of *Chromatin Organization* and *Developmental Biology* pathways are almost exclusively enriched in the networks corresponding to oligodendrocyte differentiation. Interestingly, a set of pathways linked to cell cycle like *Cyclin D associated events in G1* (R-HSA-69231) are enriched in the oligodendrocyte case study and reported to be also relevant in glioblastoma^41–44^. All case studies appear constitutively enriched in a cluster of *Pathways in Cancer* annotated downstream of *Diseases of signal transduction by growth factor receptors and second messengers* (R-HSA-5663202). We investigated this potential limitation in context-resolution and found that i) KNet-employed, binned network distances between genes in R-HSA-5663202 subpathways are not significantly smaller than those within other tested pathways; ii) Spearman correlations between p-values and network or corpus sizes are equivalent in all tested pathways; iii) R-HSA-5663202 subpathway categorization is associated with lower p-values both globally and within the same major topic (**Supp. Fig. 5**). Perhaps unsurprisingly, we concluded that proteins from these pathways like MAP-kinases and PKB are generally involved in the explored case studies; this also suggests that the observed clustering of cancer-related studies is not exclusively dependent on the enrichment of cancer pathways. Finally, we quantitatively assess the context resolution of the ENQUIRE-informed enrichment (**Fig. 5C**). To this end, we performed a permutation test on the observed Baker’s gamma correlation value between dendrograms (**Fig. 5A**-**B**), which allows to statistically assess their similarity^45^. We benchmarked its significance against two other methods, namely gene set over-representation analysis (ORA), and topology-based pathway enrichment analysis using STRING’s high-confidence functional associations, instead of ENQUIRE-generated co-occurrences, to compute the 𝑄 node scores (see **Mat.Met.**). All methods generated a dendrogram significantly closer than expected to the reference. In our analysis, topology-based enrichments outperform ORA, with the ENQUIRE-informed score moderately improving the performance over the STRING-informed equivalent (0.69 and 0.64, respectively). Taken together, these results suggest that ENQUIRE-generated networks can effectively represent contextual, biological differences and similarities between case study corpora. While ENQUIRE-annotated genes are sufficient for context resolution, the use of topology-based methods that incorporate corpus-specific co-occurrence information improves the performance.

**Fig. 5.**
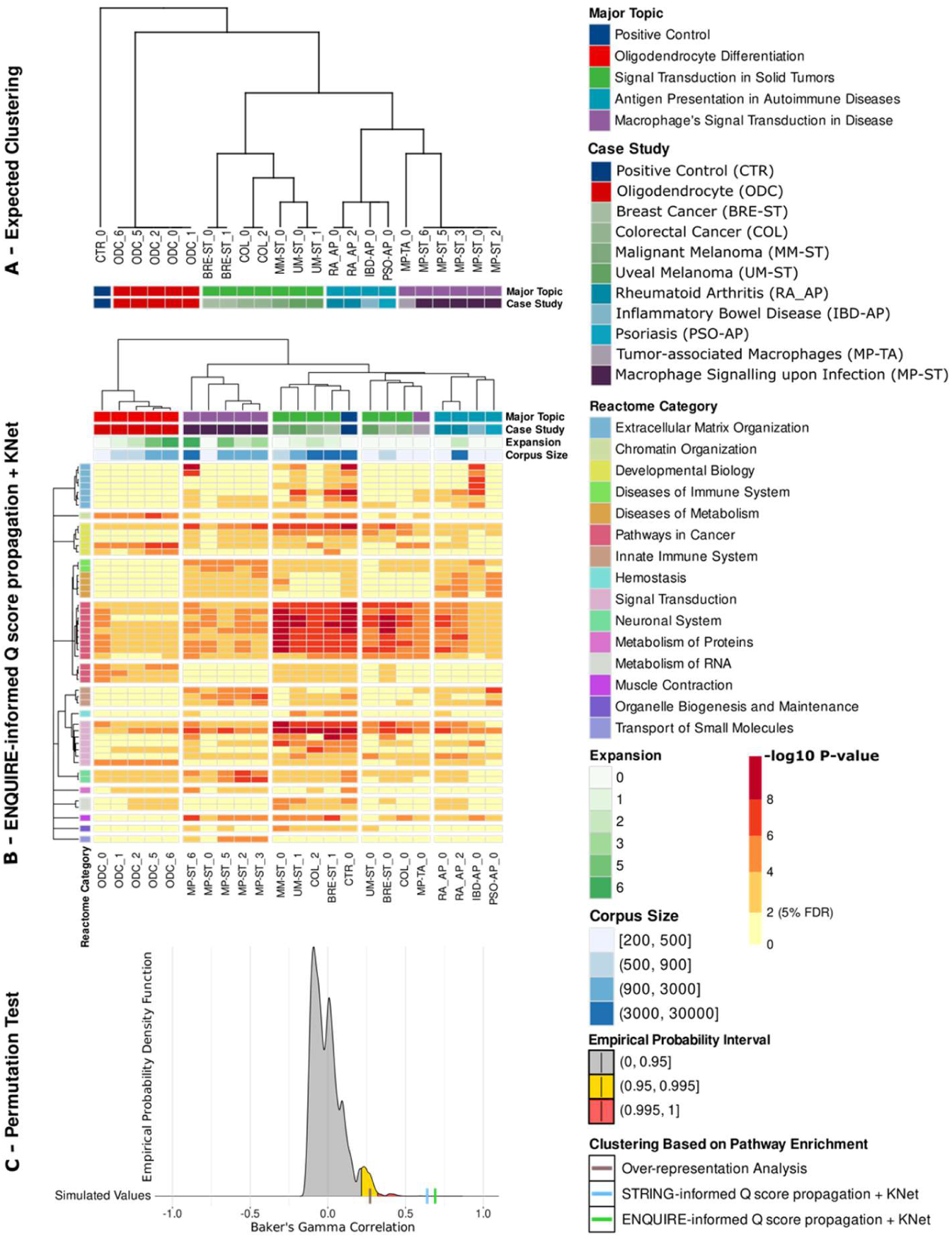
ENQUIRE-generated graphs enhance the context resolution of pathway enrichment analyses. A: reference dendrogram showcasing the expected categorization of the case studies described in **Table 1**. The number following a case study abbreviated name indicates the expansion counter. Network expansions that did not yield any new gene were excluded. B: Topology-based pathway enrichment, obtained by applying 𝑄 score propagation and SANTA’s KNet function on ENQUIRE-informed gene-gene associations (see Post Hoc Analyses under **Mat.Met.**). The heatmap shows the unadjusted p-values for the 50 enriched Reactome pathways with at least one significant, adjusted p-value (5% FDR) and highest variance across case studies (the dendrogram was computed on the complete statistic). Pathways are clustered according to Reactome’s internal hierarchy. We respectively apportioned the dendrograms into 5 and 15 partitions to visualize their coherence to Major Topic and Reactome Categories. Legends for expansions, rounded corpus size, and p-values ranges are provided. C: Permutation tests of Baker’s gamma correlation between the reference dendrogram (A) and clustering obtained from alternative pathway enrichment analyses, as in. B. Colored areas indicated probability intervals obtained from simulating correlations between reference and sampled dendrograms. See **Mat.Met.** for further details.

**Table 6.** Empirical quantiles of DeltaCon similarities, ENQUIRE- and STRING-based edges counts, sorted by number of genes in the network. . Median values with respect to each metric and range of gene counts are highlighted in bold.

| Metric | Range of gene counts | Quantiles |  |  |  |  |
| --- | --- | --- | --- | --- | --- | --- |
|  |  | 0% | 25% | 50% | 75% | 100% |
| DeltaCon | 4-9 | 0.75 | 0.83 | **0.87** | 0.94 | 1.00 |
|  | 10-14 | 0.67 | 0.78 | **0.81** | 0.83 | 1.00 |
|  | 15-23 | 0.65 | 0.74 | **0.77** | 0.79 | 0.87 |
|  | 24-119 | 0.56 | 0.65 | **0.69** | 0.72 | 0.81 |
| Edge count - ENQUIRE | 4-9 | 4 | 6 | **7** | 8 | 16 |
|  | 10-14 | 6 | 8 | **10** | 13 | 43 |
|  | 15-23 | 8 | 13 | **17** | 22 | 66 |
|  | 24-119 | 18 | 36 | **49** | 77 | 295 |
| Edge count - STRING | 4-9 | 4 | 6 | **8** | 10 | 23 |
|  | 10-14 | 6 | 11 | **15** | 20 | 50 |
|  | 15-23 | 10 | 21 | **28** | 37 | 94 |
|  | 24-119 | 19 | 54 | **89** | 146 | 591 |
| Connected components - ENQUIRE | 4-9 | 1 | 2 | **2** | 3 | 5 |
|  | 10-14 | 1 | 3 | **4** | 5 | 8 |
|  | 15-23 | 1 | 4 | **6** | 7 | 12 |
|  | 24-119 | 1 | 4 | **6** | 7 | 15 |
| Connected components - STRING | 4-9 | 1 | 1 | **2** | 2 | 5 |
|  | 10-14 | 1 | 2 | **2** | 3 | 6 |
|  | 15-23 | 1 | 2 | **3** | 4 | 8 |
|  | 24-119 | 1 | 2 | **2** | 4 | 12 |

## DISCUSSION

ENQUIRE is a novel computational framework that combines textmining, network reconstruction, and literature querying, offering an alternative to manual literature curation and database mining. ENQUIRE interrelates gene mentions and biomedical concepts through co-occurrence networks and tabulated references while accounting for biases in the input literature corpus. Its framework enables *post hoc* analyses that infer contextual gene sets and enriched molecular pathways. ENQUIRE can enhance the biological interpretation of omics data, suggest relevant processes and components for computational models, and motivate the selection of molecular targets for biological experiments and in scenarios like molecular tumor boards. We opted for a compromise between coverage of unannotated article abstracts (gene normalization) and high-fidelity, pre-computed concept annotations (MeSH retrieval). ENQUIRE’s gene normalization strategy is appropriate for reconstructing co-occurrence gene networks with affordable computational requirements, and scales well with large input corpora, without the need of restricting the analysis to databases of pre-annotated gene mentions^46^. The combination of a curated lookup table with abstract-specific blocklists enhances precision, thus leading to co-occurrence networks with fewer false positives, compared to recall-oriented approaches like BERN2^35,47^. An added value of ENQUIRE is that the obtained gene/MeSH co-occurrence network can prime further information retrieval beyond textmining. Differently from previous works on gene/MeSH relations, our statistical framework is independent of the user scope (genes or MeSH can be mined separately) and is not immutable with respect to a species or general topic (e.g. diseases)^48–51^. Instead, ENQUIRE automatically constructs PubMed queries from network-derived genes and MeSH to expand the input corpus, and in turn the network. We also assessed ENQUIRE’s performance using real-world case scenarios. For example, we investigated the relationship between ENQUIRE-suggested co-occurrences and database-annotated gene interactions. Our results indicate that ENQUIRE-generated gene co-occurrence networks reflect experimental and database-annotated functional gene associations. At the same time, ENQUIRE can also generate networks with previously unannotated wirings that can encourage novel explorative analyses (**Fig. 4B**). We also analyzed the feasibility of corroborating ENQUIRE-suggested relations by mapping co-occurrence information onto a mechanistic reference network. Since there is no generalizable method to project a network of indirect relations (co-occurrences) onto a mechanistic network^52–56^, we designed a function to score a physical interaction network using ENQUIRE-generated networks. This allowed us to verify that the enriched pathways in original and expanded ENQUIRE networks reflect their contexts and enable the comparison of multiple case studies. This strategy still poses some limitations in terms of choosing a reference network and pathways to be tested. We designed ENQUIRE as a series of modular, open-source components that can be combined and expanded to tune its performance. For instance, one could insert a part-of-speech recognition parser upstream of the co-occurrence detection step to strengthen its criteria^57^. Similarly, one can implement a propensity matrix into the random graph model to further weight a co-occurrence with its textual context^14,21^. As gene normalization relies on the utilized lookup table of reference gene symbols and aliases, ENQUIRE’s accuracy depends on how comprehensive and free of ambiguities this table is. The current version of our algorithm only performs normalization of human genes and corresponding mouse orthologs. Still, it can be adapted to perform gene normalization of any other species by supplying an appropriate lookup table, such as those provided by the STRING database^58^. Our main objective was to construct a robust textmining, network reconstruction, and automatic querying pipeline accessible to bioinformaticians and systems biologists with affordable computational requirements. Since the standalone version of the algorithm requires some background in computer programming, we are working to provide a web version of ENQUIRE to ease its adoption among biomedical researchers.

## DATA AVAILABILITY

ENQUIRE’s main program and the standalone scripts to perform the *post hoc* analyses are included in an Apptainer/Singularity image file (SIF), available for download at https://figshare.com/articles/software/ENQUIRE/24434845 (DOI: 10.6084/m9.figshare.24434845.v3). Installation and running instructions, gene-symbol-to-alias lookup table, input and output files from the exemplary case study, and data underlying the results (**Supp. Information**) can be found at https://github.com/Muszeb/ENQUIRE (DOI: 10.5281/zenodo.10692274). All the individual scripts are also available upon request.

## AUTHOR CONTRIBUTIONS

Idea and concept: LM and JV. Coding and benchmarking of the algorithm: LM and MW. Drafting of the manuscript: LM, XL, and JV. All the authors edited, corrected, and approved the submitted draft.

## Supporting information

Data Availability Statement

## ACKNOWLEDGEMENTS

We thank Martin Eberhardt, Christopher Lischer, Jimmy Retzlaff, Esther Güse, and Suryadipto Sarkar for the useful scientific discussions, comments on the manuscript, and testing the installation and running of the algorithm.

## MATERIALS AND METHODS

### Description of the ENQUIRE algorithm

#### Extraction of Article Metadata

ENQUIRE uses the NCBI’s e-utilities to query and fetch information from the PubMed database^59^. *Epost* is used to request a collection of PMIDs, *efetch* to extract their metadata in XML format, and *esearch* to construct PubMed queries.

#### MeSH Term and Article Abstract Extraction

For each MEDLINE-indexed, input PMID, if the MeSH entity scope is selected, ENQUIRE retrieves MeSH main headings (“descriptors”) and subheadings (“qualifiers”) from their respective *efetch*- retrieved XML files. These MeSH terms are further selected to match biomedically relevant, non-redundant categories, by exploiting the tree-like, hierarchical structure of the MeSH vocabulary. By default, ENQUIRE only retains members downstream of the MeSH categories A (Anatomy), C (Diseases), D (Chemicals and Drugs), and G (Phenomena and Processes), except for sub-categories G01 (Physical Phenomena), G02 (Chemical Phenomena) and G17 (Mathematical Concepts).

#### Gene Normalization from Article Abstracts

For each input PMID, if the gene entity scope is selected, ENQUIRE retrieves article abstracts from their respective *efetch*-retrieved XML files. As other authors have shown that the proportion of gene mentions does not significantly differ between abstracts and full-body texts^60^, we only mine the abstracts for gene mentions. In contrast to standard named entity recognition of genes (NER), whose task is to exactly match the character span of a gene mention, ENQUIRE’s textmining framework aims at detecting least one gene alias per unique reference gene mentioned in an abstract. We therefore designed a “Swiss cheese model” for gene normalization, in which multiple methods complement each other to improve the global precision. In brief, ENQUIRE applies up to two algorithms to each unprocessed abstract: i) the Schwartz-Hearst algorithm to detect single-word abbreviations and their respective definitions^33^; ii) the optional scispaCy model (*en_ner_jnlpba_md*) to identify words classified as “CELL_LINE” or “CELL_TYPE”^34^. This allows ENQUIRE to construct abstract-specific blocklists that discard i) ambiguous abbreviations whose definitions are not similar to any gene alias from a pre-annotated lookup table, and ii) ambiguous or unwanted mentions to cell entities containing gene aliases, such as “CD8+ T cell”. Finally, a tokenization module generates potential gene-alias-matching tokens and redirects them to a unique, reference gene symbol using the lookup table.

#### Construction of the Lookup Table of Reference Gene Names and Respective Aliases

Similar to previous approaches^61^, ENQUIRE performs NER of *Homo sapiens* and *Mus musculus* gene mentions, while also redirecting the latter to their respective human homologues using MGI’s mouse/human orthology table^62^. Each reference gene name corresponds to HGNC approved symbol^63^. Additional mouse and human gene aliases were pooled from HGNC (“previous symbols”, “previous names”, “alias symbols”, “alias names”), ENSEMBL (“gene stable ID”, “gene description”, “gene name”), Uniprot (“gene names”, “protein names”), and miRBase (“ID”, “alias”, “name”)^64–66^. We manually inspected sources of ambiguities and lack of spelling variants: for example, we added miRNA names without species suffixes (e.g. “miR-335” from “hsa-miR-335”), multiple spellings for lnc- and mi-RNAs (e.g. “LNC/Lnc/lnc”, “miR/mir”) and removed aliases identical to common acronyms for experimental techniques (e.g. “MRI”, “NMR”, “TEM”). We converted Greek letters to their literal spelling. We resolved ambiguities due to aliases reported under more than one reference symbol, by either assigning the alias to a single reference, or by excluding the alias.

#### Abstract Tokenization for Named-Entity Recognition of Genes

ENQUIRE mostly performs named-entity recognition of genes (NER) from article abstracts by exact matches between gene aliases and space- or punctuation-separated word tokens. We exclude general-purpose English words annotated in the *English-words* Python library to reduce the computational burden of mapping gene mentions. Greek letters are converted to their literal spelling. Special attention is put to hyphen- and slash-containing tokens, tracing their usage as integral parts of gene aliases (e.g. “TNF-alpha”) or separators (e.g. “FcγR-TLR Cross-Talk” – PMID 31024565, “Akt/PI3K/mTOR signaling pathway” – PMID 35802302). When cases of the latter kind occur, the algorithm requires all hyphen- or slash-separated words to be gene aliases, in order to be considered individual tokens. Then, ENQUIRE tokenizes the abstract into single-word tokens and interprets unambiguous tokens as the corresponding reference gene symbol if they match an alias in the lookup table. Multiple mentions of the same gene within an abstract count as one.

#### Abstract-Specific Blocklists Using Cell Entity Mentions and Abbreviation-Definition Pairs

Any token that exactly matches an alias from the lookup table is redirected to the respective reference symbol, except when that same token is either classified as part of “CELL_LINE” or “CELL_TYPE” entities, or as an abbreviation, by scispaCy *en_ner_jnlpba_md* and Schwartz-Hearst models. In the former exception, the token is added to a blocklist and any of its mentions within the abstract text are excluded from further gene normalization steps. In the latter exception, we evaluate the validity of an alias-matching abbreviation by means of its definition, as inferred by Schwartz-Hearst. We perform string comparison to calculate alignment scores between the definition and any recorded alias of the same reference symbol matched by the abbreviation. To this end, we implemented the Needleman-Wunsch algorithm for global alignment, with match score equal to 1, gap opening and mismatch penalties equal to -1, and gap extension penalty equal to -0.5^67^. Next, we calibrated a threshold for either retaining or discarding an alias-matching abbreviation according to its optimal alignment score. We used a dataset of abbreviation-description pairs from more than 300 abstracts and generated a distribution of scores by aligning any description to any annotated alias. Intuitively, there could only be a handful of alignments between an actual gene description and the aliases referring to that same gene, as opposed to several alignments between that same description and unrelated aliases. Therefore, we treated the above derived distribution as a model describing false positive alignments between descriptions and gene aliases. Finally, we identified a range between 0.1 and 0.2 that respectively correspond to 95th and 99th percentiles of the distribution of alignment scores as a sensible interval for choosing the threshold. We opted for a threshold of 0.15. Therefore, for any description whose abbreviation matches a gene alias, ENQUIRE records a gene mention only if the maximal alignment score against any alias of that same gene is higher or equal to this threshold; else, the abbreviation is added to the blocklist and all of its mentions within the text are excluded. Notice that the blocklist is independently computed for each abstract, thus making ENQUIRE’s gene normalization moderately adaptive with respect to syntactical context.

#### Annotation and Weighting of Co-Occurrences

ENQUIRE records the occurrences of MeSH and gene entities within each input article. Then, it counts pairwise co-occurrences by enumerating the subset of PMIDs associated to both entities in each pair. For each pair of entities 𝑔_*i*_and 𝑔_*j*_that co-occur in at least one article, we define the weights 𝑤 and distances 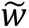 accounting for the sheer co-occurrence 𝑋(𝑔_*i*_, 𝑔_*j*_) as follows:

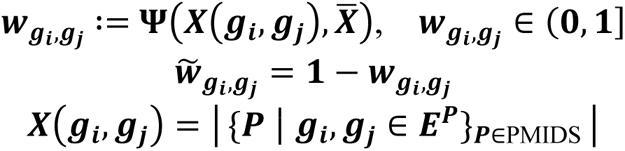

Where *X̄* is the mean co-occurrence between any two entities in the corpus, Ψ(⋅, *X̄*) is the zero-truncated, Poisson cumulative density function with a lambda of *X̄*, and 𝐸^*P*^ is the set of all entities annotated within the PMID 𝑃 that belongs to the submitted PMIDS corpus. This scoring system assigns higher relevance to co-occurrences that appear more often than average.

#### Reconstruction of a Weighted Network of Significant Co-Occurrences

ENQUIRE converts the recorded co-occurrences into an undirected multi-graph, where gene or MeSH terms become nodes, and each recorded co-occurrence between two entities becomes an edge. Thus, the network has as many nodes as the number of unique MeSH and gene symbols, with as many edges between two nodes as the number of PMIDs in which they co-occur. ENQUIRE implements the Casiraghi-Nanumyan’s soft-configuration model applied to undirected, unweighted edge counts to select significant co-occurrences among entities, adjusted to 1% FDR^21^. The test statistics follows a multivariate hypergeometric distribution, under the null hypothesis of observing a random graph whose expected degree sequence correspond to the observed one. This allows us to condition the testing to the sheer, per-entity occurrence, which serves as a proxy for leveraging literature biases in the corpus. It is important to note that the null model does not assume independence of individual edges, but merely their equiprobability, and is unaffected by the weights w. This selection results in an undirected, single node-to-node edge co-occurrence graph (i.e. a simple graph). For each pair of adjacent entities g_i_ and g_j_ in the simple network, we assign the weights w_gi, gj_ and distances 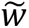_gi, gj_ to their mutual edge. Additionally, we prune poorly connected nodes by modularity-based, w-weighted Leiden clustering^68^ and removal of communities that consist of a single node. From the resulting gene/MeSH network, we also extract the respective gene- and MeSH-only subnetworks.

ENQUIRE-generated gene/MeSH networks can consist of multiple connected components, i.e. subgraphs. To exclude unimportant components, a subgraph 𝑆 is retained for subsequent computations only if the fraction of corpus articles covered by 𝑆 is higher than a threshold value, as formally defined in

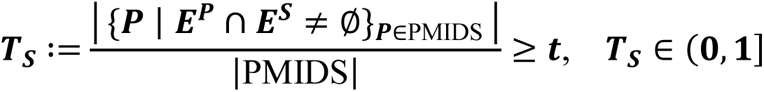

where 𝑃 denotes a PMID belonging to PMIDS, and 𝐸^*P*^and 𝐸^*S*^refer to the sets of gene or MeSH entities recorded in either 𝑃 or 𝑆. Therefore, 𝑇_*S*_ reflects the representativeness of 𝑆 with respect to the entirety of the submitted corpus. The value of 𝑡 can be set by the user. To avoid introducing irrelevant entities, ENQUIRE stops without further network expansion if the gene/MeSH network and the respective gene- and MeSH-only subnetworks individually contain only a single, connected component with 𝑇_*S*_ ≥ 𝑡. We compute the weight of a node 𝑔 in the connected graph 𝑆 utilizing the composite function 𝑊, which is the product of normalized metrics for betweenness centrality (𝑏) and 𝑤*-*weighted degree strength (𝑑):

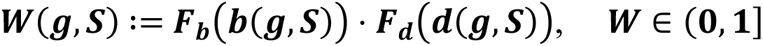

Here, 𝐹_*x*_ denotes the empirical cumulative density function for the corresponding 𝒙 parameter, calculated over 𝑆.

## Construction of Communities from “Information-Dense” Cliques

To identify the most relevant parts of the gene/MeSH network, ENQUIRE first identifies the maximal cliques of order three or more. By definition, these are graphlets whose nodes are all adjacent to each other and not a subset of a larger clique. Applying the KNet function from the SANTA R package^40^ to the gene/MeSH network having distances 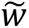_gi, gj_, we select cliques that form significant clusters of associated entities. The permutation test procedure internal to KNet allows us to consider the network topology and adjust each maximal clique’s significance, in case many other cliques of similar size exist in the network. We set the significance level for this test to 1% FDR. Subsequently, ENQUIRE generates a pruned network 𝐶 containing only statistically significant cliques. Here, ENQUIRE stops if the gene/MeSH network contains less than two significant cliques according to KNet. Next, ENQUIRE identifies communities in the 𝐶 network using modularity-based, 𝑤 -weighted Leiden clustering. ENQUIRE stops if it detects a single community that encompasses all nodes in 𝐶.

## Identification of Community-Connecting Entities

For any two disjoint communities 𝐶_*i*_ and 𝐶_*j*_, we select the set of community-connecting, weighted graphlets Γ*_C_i_,C_j__*(𝑉_*k*_, 𝐿_*k*–1_) satisfying the properties: i) all nodes 𝑔_i_ in the 𝑘-sized set 𝑉_*k*_ belong to either 𝐶_*i*_ or 𝐶_*j*_; ii) the intersections between 𝑉_*k*_ and 𝐶_*i*_ or 𝐶_*j*_ are non-empty; iii) the 𝑤-weighted, 𝑘 − 1 edges 𝐿_*k*–1_ are sufficient to obtain a single connected component; iv) there is only one edge 𝑙_gi,gj_ that connects nodes belonging to distinct communities. Here, 𝑘 is a parameter chosen by the user.

This allows us to rank the set of community-connecting entities 𝑉_*k*_in any graphlet Γ*_C_i_,C_j__* by means of the distance metric 𝑅:

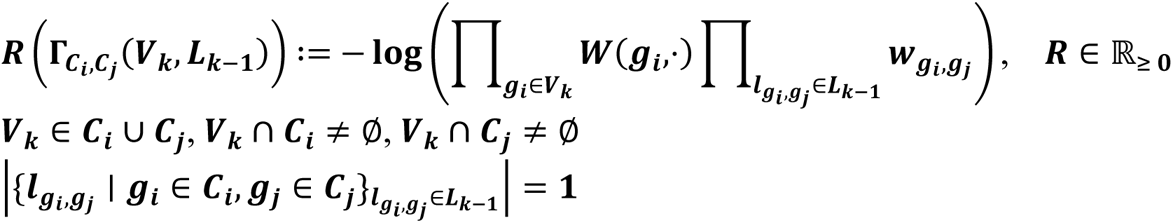

The smaller 𝑅, the closer two communities connected by 𝑉_*k*_ are.

### Retrieval of New PMIDs via PubMed Queries Based on Optimal Connections

To evaluate which genes and MeSH terms are particularly suited for expansion querying, ENQUIRE constructs a multigraph 𝑀 where network communities become nodes and all 𝑅-weighted connections between two communities become edges. 𝑅-weighted edges that do not fulfil the triangle inequality 𝑅 (Γ*_C_i_,C_j__*) ≤ 𝑅(Γ*_C_i_,C_z__*) + 𝑅 (Γ*_C_z_,C_j__*) , ∀ 𝑖, 𝑗, 𝑧 are excluded. Then, we solve the travelling salesman problem (TSP) utilizing Christofides’ approximate solution as implemented in the Python package Networkx^69^. Via the visited edges, this yields an optimal path across communities and a corresponding collection of 𝑉_*k*_entity sets. Each selected 𝑘-sized set 𝑉_*k*_ results in a PubMed query formulated via the NCBI’s *esearch* utility^59^. We condition the search terms representing gene aliases and MeSH with “[Title/Abstract]” and “[MeSH Terms]”, respectively, and exclude review articles from the results. The constructed PubMed queries require a match for all the 𝑘 entities in the optimal path – e.g. *“melanoma/immunology”[MeSH Terms] AND (“IL1B”[Title/Abstract] OR “interleukin 1-beta”[Title/Abstract] […]) AND […]*. If all queries involving a subset of the network communities lead to empty results, we prune all previously used edges from 𝑀, compute a new TSP solution, and submit newly generated queries, provided at least one entity per query belongs to such community subset. This process is repeated 𝐴 times, where 𝐴 is a parameter specified by the user. If at least 1 new PMID matches any of the constructed queries, ENQUIRE starts a new analysis from the union of new and old PMIDs; otherwise, it stops. The rationale behind merging old and new PMIDs is to account for the original corpus when computing the statistics on new co-occurrences.

#### Post-hoc Analyses

### Context-Aware Gene Sets

To reconstruct contextual gene sets using gene/MeSH co-occurrence networks, we adapt network-based relational data to the method described by Khan *et al.*^70^. To this end, we first construct the inverse log-weighted similarity matrix between the gene/MeSH network nodes^71^. This metric prioritizes nodes sharing many lower degree neighbors rather than few higher degree ones. We derive a Euclidean distance matrix from the similarity matrix, after applying a Z-score standardization; then, we use the R package DynamicTreeCut and Ward’s clustering to identify initial clusters and create an initial membership degree matrix^72,73^. Finally, we detect fuzzy clusters of genes and MeSH terms by applying Fuzzy C-means clustering to the Euclidean distance matrix, using the R package ppclust^1,2^. The resulting membership degree matrix allows annotating genes with desired cluster membership degrees and extracting the linked MeSH terms to characterize the gene set.

### Context-Aware Pathway Enrichment Analysis

We designed a method to map any text-mined co-occurrence network G onto a mechanistic reference network 𝑁 and infer context-specific enrichment of molecular pathways. With this strategy, we attempt to mechanistically explain the indirect relationships that constitute the co-occurrence network. To this end, we define the fitness score 𝑄 for every gene 𝑔 in 𝑁 with non-zero node degree 𝑑:

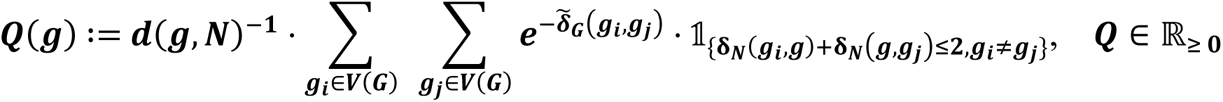

Here, 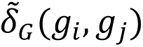 and 𝛿_*N*_ (𝑔_i_, 𝑔_j_ ) are the 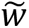-weighted and unweighted distances from 𝑔*_i_* to 𝑔*_j_* in the graphs 𝐺 and 𝑁, respectively. The indicator function 𝟙 implies that non-text-mined genes without at least two text-mined nodes as neighbors have 𝑄 equal to zero. We normalize all scores to decorrelate Q from the node degree 𝑑, similarly to other approaches in network propagation^74,75^. As a mechanistic reference network, we chose STRING’s (release 11.5) *H. sapiens* network of protein-coding, physically interacting genes^38^. We exclusively combined the “experimental” and “database” channels to calculate STRING’s confidence score, then pruned all edges with score below the 90^th^ percentile. After removing zero-degree nodes, we obtain a reference, unweighted network of 9,482 nodes and 88,333 edges. Then, we calculate 𝑄 scores for protein-coding genes in the STRING reference network (𝑁), using the ENQUIRE-generated gene network (G). We test for associations between predefined gene sets and high-scoring node clusters using SANTA’s KNet function^40^. KNet takes as input the STRING reference network, its nodes’ 𝑄 scores, and a gene set; it then tests if the latter is enriched, based on scores and graph distances of protein-coding genes belonging to both the network and the gene set. This way, we aim at capturing known experimentally or database-derived molecular interactions relevant to ENQUIRE’s input literature corpus, using topology-based enrichment analysis. We test for enrichment on gene sets derived from Reactome pathways, obtained via the Reactome Graph database^37^. See **Supp. Fig. 2** for an example of 𝑄 score weighting.

## Benchmarks and Case Studies

### Assessment of ENQUIRE’s Gene Normalization Accuracy and Performance

We evaluated ENQUIRE’s gene normalization precision and recall using abstracts from the NLM-Gene corpus mentioning at least one *M. musculus* or *H. sapiens* gene – 479 out of 550 entries^32^. We tested the four module combinations obtained by either including or excluding the cell entity recognition module *en_ner_jnlpba_md* and the *Schwartz-Hearst* abbreviation-definition algorithm^33,34^. We compared the computational performance of ENQUIRE’s gene normalization method using both *en_ner_jnlpba_md* and *Schwartz-Hearst* against GNorm2 implementation of Bioformer^36,35^. We computed wall time by accounting for both text processing and loading of required data such as gene alias lookup tables and machine learning models. RAM usage was measured using resident set size (RSS) measurements returned by the Linux built-in function *ps*. We ran the computations on a Linux computer with 20 CPUs (3.1 GHz) and 252 GB of RAM. Up to 8 cores were used for parallelization.

### Inference of Reactome Gene Sets from Reference Literature

We extracted annotated genes and reference literature for all *H. sapiens* Reactome pathways from the Reactome Graph database^37^. We employed NCBI’s *esearch* and e*link* utilities to retrieve primary research articles cited by review articles^59^. After excluding pathways with less than three primary literature references or only one annotated human gene, we obtained a set of 967 pathways. For each pathway literature corpus, ENQUIRE performed one network reconstruction, set to only extract gene mentions from article abstracts. We evaluated the effects of corpus size, pathway size, and average gene-gene co-occurrence per abstract on precision and recall of ENQUIRE’s gene normalization and network reconstruction. We also evaluated the correlation between true positives and the corpus- and network-based node weight 𝑊.

### Estimate of Molecular Interrelations

We automatically generated a list of case studies by crossing leaf nodes downstream of *Diseases* and *Genetic Phenomena* (G05) MeSH categories. We then constructed a PubMed query from each pair by “AND” concatenation. Examples of such queries are *“Stomach Neoplasm”[MeSH Terms] AND “Chromosomes, human, pair 18”[MeSH Terms]*, and *“Acquired immunodeficiency syndrome”[MeSH Terms] AND “Polymorphism, single nucleotide”[MeSH Terms]*. For each query result with a size between 50 and 500 articles, we executed one network reconstruction. If obtaining a gene-gene co-occurrence network, we investigated whether its set of genes produced a network with more functional interactions than expected by chance. To obtain background distributions of edge counts for each gene set size observed with ENQUIRE, we sampled one million random gene sets and cumulated their interconnecting edges in STRING’s v. 11.5 *H. sapiens* functional protein network. We only included functional associations from experiments, co-expression, and third-party databases with a cumulative score higher than 0.7 between proteins. The significance of each ENQUIRE-generated gene set’s edge count was computed from the right-tailed probability of the empirical distribution.

Moreover, we compared the ENQUIRE-generated gene-gene wirings to STRING-derived associations using the DeltaCon similarity measure in a permutation test^39^. To this end, we generated 10,000 random graphs for each observed ENQUIRE network. Each random graph was obtained through 300 random edge-swapping attempts while preserving the degree sequence of the original network. To obtain sensible probability densities, we focused on ENQUIRE-generated networks with degree sequences allowing at least ten different realizations of a graph. We followed the formula 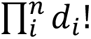, where 𝑑_*i*_ is the degree of the 𝑖-th node of a graph containing 𝑛 nodes.

### Assessment of Context Resolution by Topology-Based Enrichment of Molecular Pathways

To show that ENQUIRE preserves context-specific molecular signatures, we designed a broad panel of case studies (**Table 1**). Each corpus consisted of the union of references contained in three independent reviews accessible via NCBI’s *elink* utility^59^. We selected reviews from the results of PubMed search queries consisting of two or three MeSH terms (e.g. *“Melanoma”[MeSH Terms] AND “Signal Transduction”[MeSH Terms]*), favoring PubMed-ranked best matches when possible. We also included an unspecific positive control group consisting of the union of all context-specific corpora. This experimental design allowed us to construct a reference dendrogram that clusters the case studies only based on baseline biological knowledge, expecting expanded networks of a case study to cluster together with the originally reconstructed one. Then, we applied ENQUIRE with default parameters to each case study and analyzed all resulting gene-gene networks, i.e., from original and expanded corpora. We computed pairwise similarities between node and edge sets of the constructed networks using Szymkiewicz-Simpson overlap coefficient (OC):

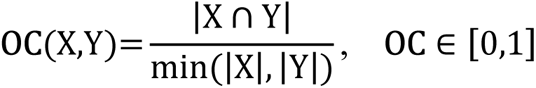

Where 𝑋 and 𝑌 are either two node sets or two edge sets. An OC of 0 indicates no overlap, while an OC of 1 indicates the smaller node or edge set is a subset of the larger one. By construction, same-case-study original and expanded networks possess OCs of 1 with each other. We applied the *post hoc*, context-aware pathway enrichment analysis described above to all generated networks. We tested the enrichment of Reactome pathways with sizes ranging from 3 to 100 genes, categorized as in the database’s *Top-Level Pathways* and disease ontologies^37^. We performed hierarchical clustering of the networks using Euclidean distance and Kendall’s correlation based on network-specific, KNet-generated p-values. We compared the resulting dendrogram to the expected one by a permutation test of Baker’s gamma correlation using one million permutations of the original dendrogram^45^. We also compared the results to two alternative statistics: i) over-representation analysis of nodes from the ENQUIRE-generated networks (the collection of all genes observed in any case study was used as the “universe”); ii) KNet statistics, using 𝑄 scores based on STRING’s high-confidence functional association network (described above) and ENQUIRE-derived gene nodes.

## EXTENDED DATA (SUPPLEMENTARY FIGURES)

**Supplementary Figure 1.**
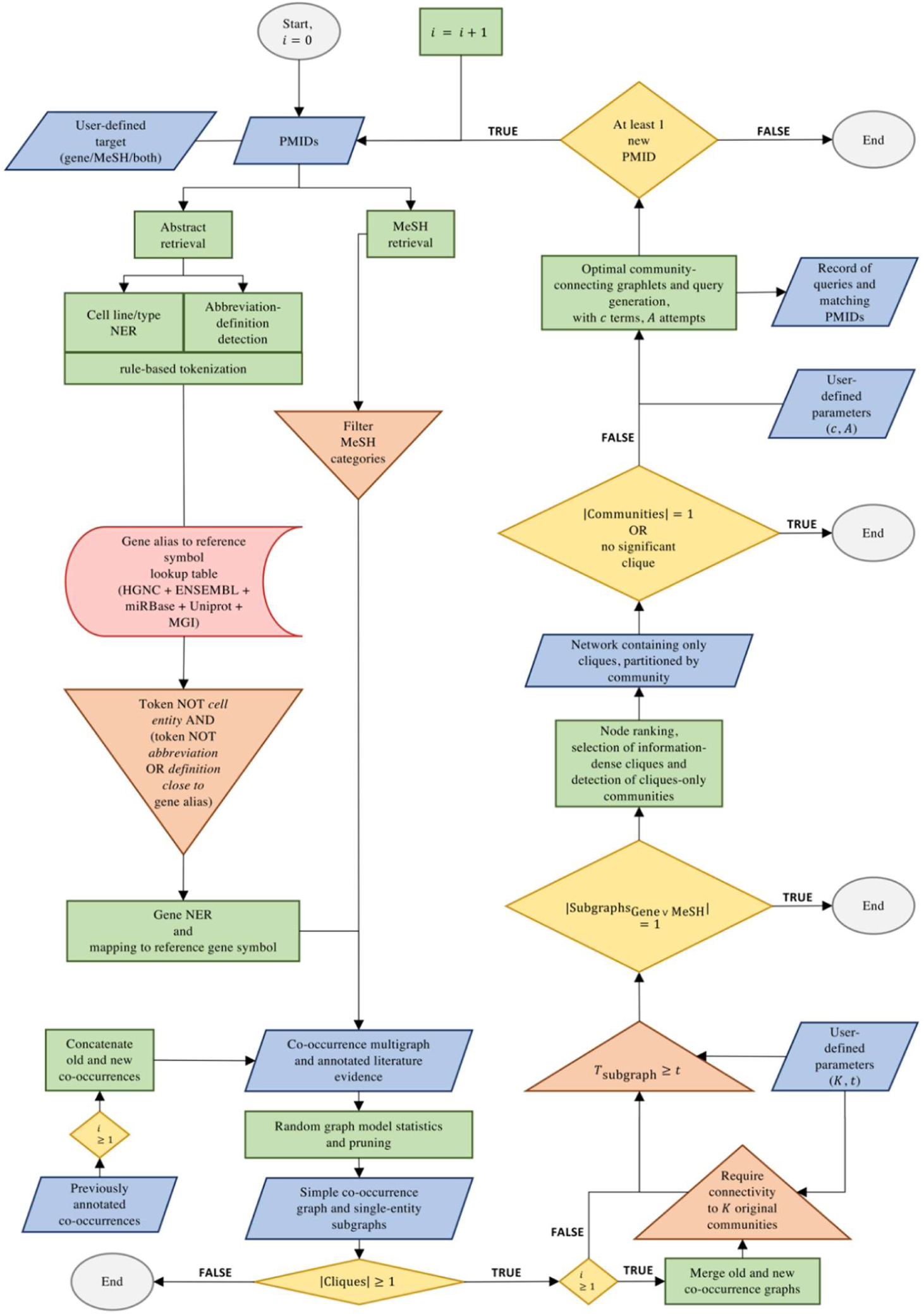
ENQUIRE’s flowchart. The pipeline’s schematics is described with respect to start and end points (grey ellipses), input, parameters, and generated data (blue parallelograms), algorithms (green rectangles), filtering (red triangles), pre-computed data (pink halfpipes), and branching points (yellow diamonds). NER: named-entity recognition. PMID: PubMed identifier. MeSH: Medical Subject Heading. Detailed explanation of the parameters and algorithms is provided in the main text.

**Supplementary Figure 2.**
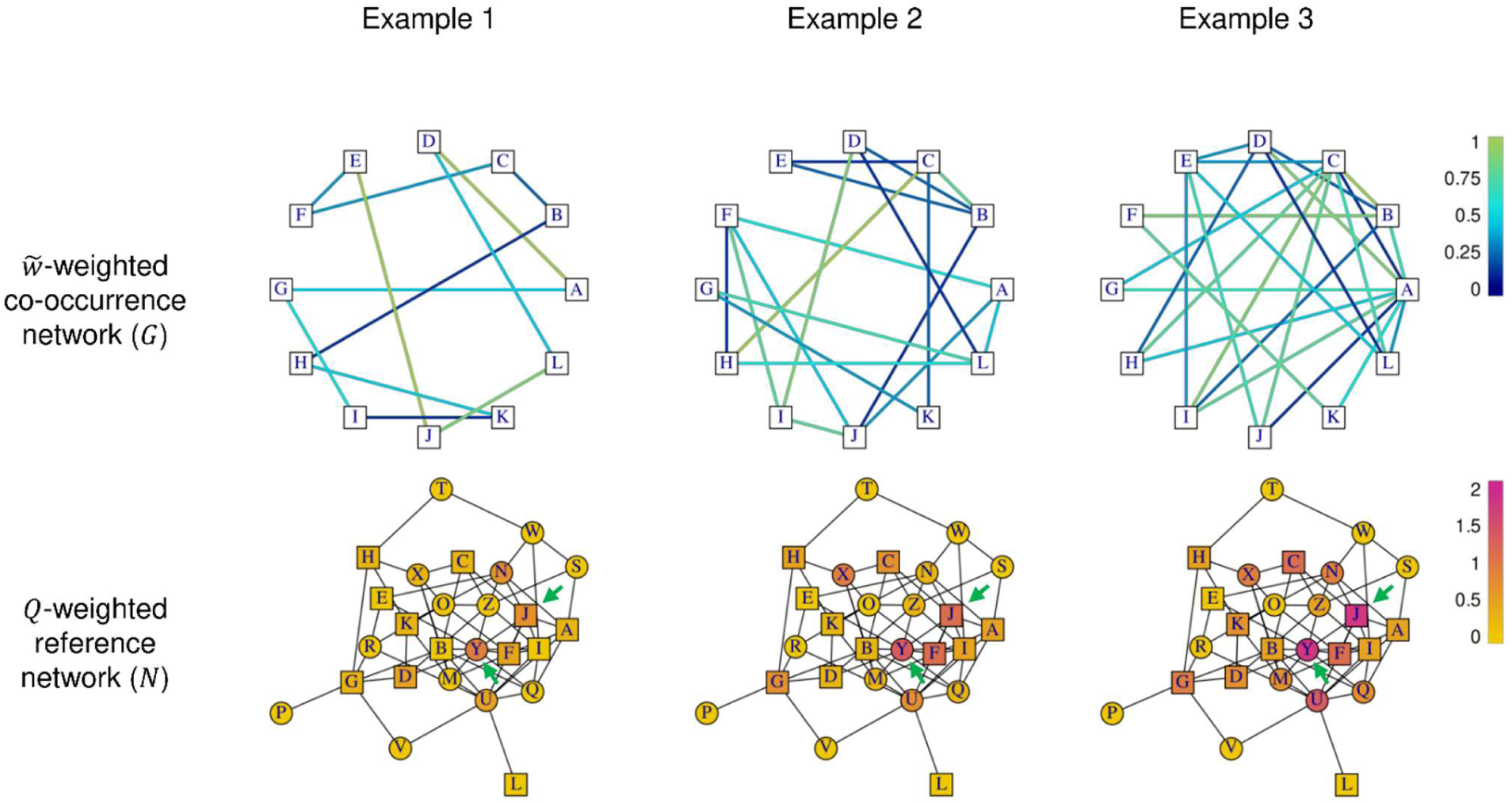
Example of Q score weighting. The top row shows three simulated co-occurrence networks 𝐺 with the same set of textmined genes (squares), generated with progressively higher edge-forming probability, and sampling edge weights 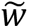 from a uniform distribution in [0,1]. Genes from an immutable reference network 𝑁 containing both textmined and non-textmined genes (circles) are weighted by the 𝑄 score. For each gene 𝑔 in 𝑁, its weight 𝑄 is a function of the textmined genes in the 𝑔-neighbourghood and their 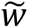-weighted distances in the network 𝐺. Nodes with relatively more connections to textmined nodes in the reference network possess higher 𝑄 scores, irrespective of being textmined or having a high node degree. See the non-textmined node Y and the textmined node J as an example.

**Supplementary Figure 3.**
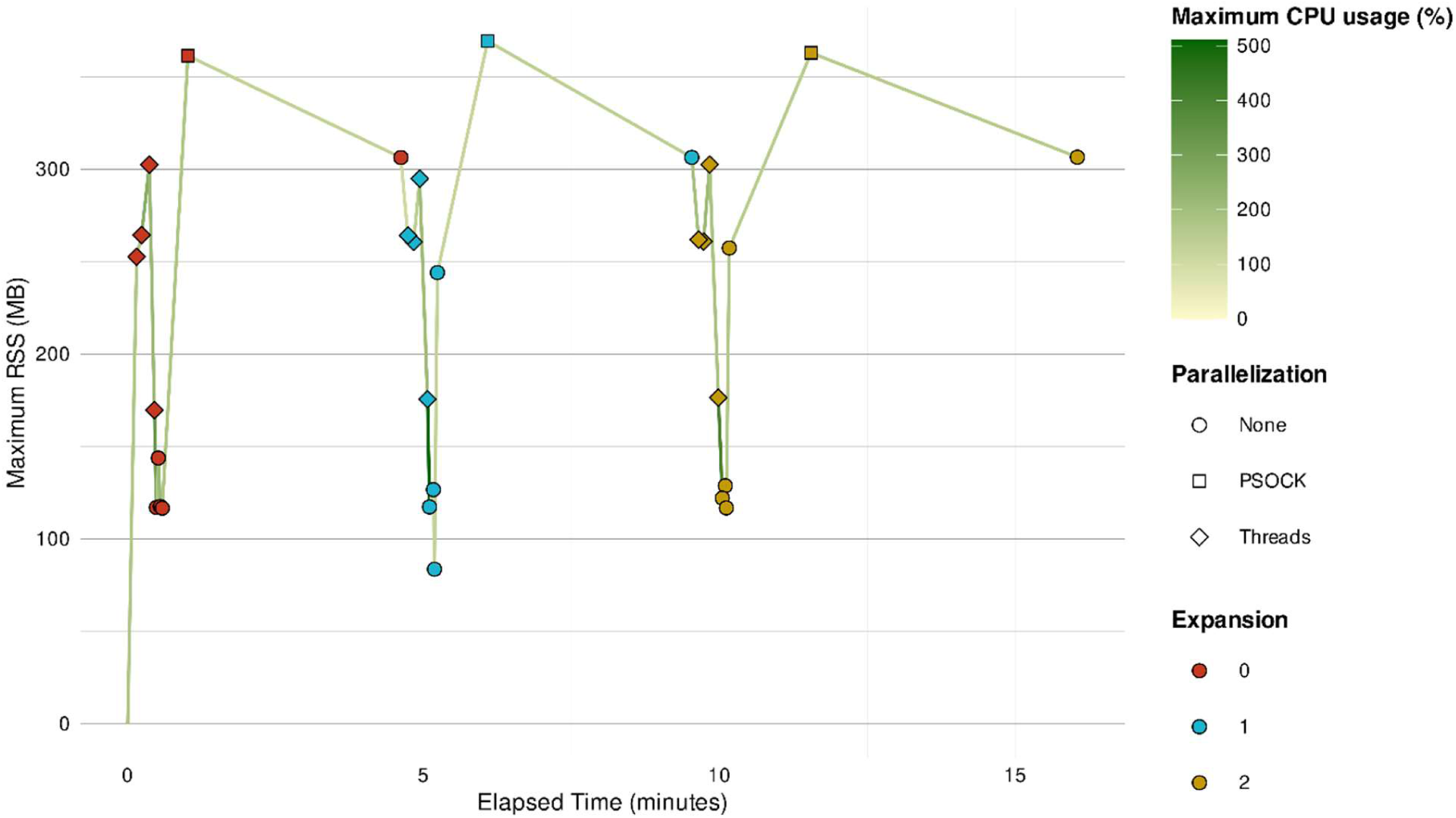
Memory and CPU usage of a typical ENQUIRE run. The chart shows the performance monitoring of the exemplary ENQUIRE run described in Results and **Fig. 2**, in which 2 expansions for a total of three iterations were performed. We used a Linux computer with 8 CPUs (2.5 GHz) and 16 GB of RAM. 6 cores were used for parallelization. Each dot represents a submodule launched by ENQUIRE, with the elapsed time at which it terminated as x-coordinate, and the maximum registered RAM usage, in the form of Resident Set Size (RSS, in megabytes), as y-coordinate. Cumulative elapsed time at the end of each reconstruction-expansion cycle is indicated. Lines in-between processes are colored by the maximum CPU usage, which is defined as the used CPU time divided by the time the process has been running, in percentage. This estimate does not typically add up to 100%. Higher CPU usage imply higher workload for each of the utilized cores. Resource usage of parallel socket cluster (PSOCK) protocol can be underestimated, as this protocol generates parallel processes whose process identifiers (PIDs) are independent of ENQUIRE’s PID and not monitored. Nevertheless, ENQUIRE restricts the memory usage of PSOCK-based parallel processes, so that their aggregated memory usage is always less than 25% of the available RAM at a given time, possibly reducing the effective number of cores used.

**Supplementary Figure 4.**
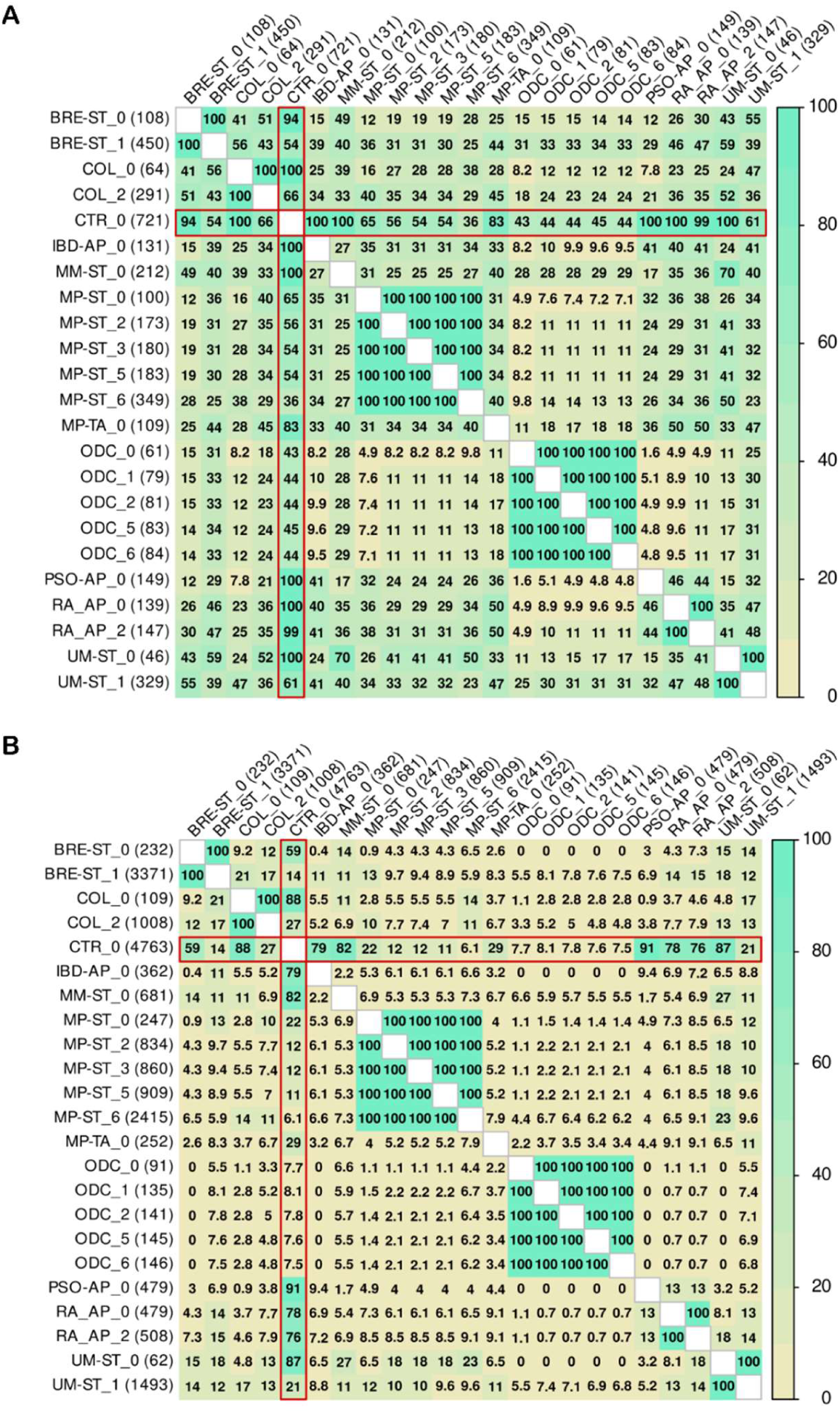
Diversity in nodes and edges from reconstructed and expanded networks generated by ENQUIRE. We computed similarity measures between ENQUIRE-inferred, co-occurrence gene networks based on the case studies described in **Tsable 1**. The number following a case study abbreviated name indicates the expansion counter. Network expansions that did not yield any new gene were excluded. Panel **A** depicts similarities between the networks’ node sets, while panel **B** depicts similarities between edge sets. Numbers and color gradient report Szymkiewicz-Simpson overlap coefficient percentages (OC). An OC of 0 % indicates no overlap, while an OC of 100% indicates the smaller node or edge set is a subset of the larger one. By construction, same-case-study original and expanded networks possess OCs of 100% with each other. OC between the positive control (CTR) and other case study networks are highlighted in red

**Supplementary Figure 5.**
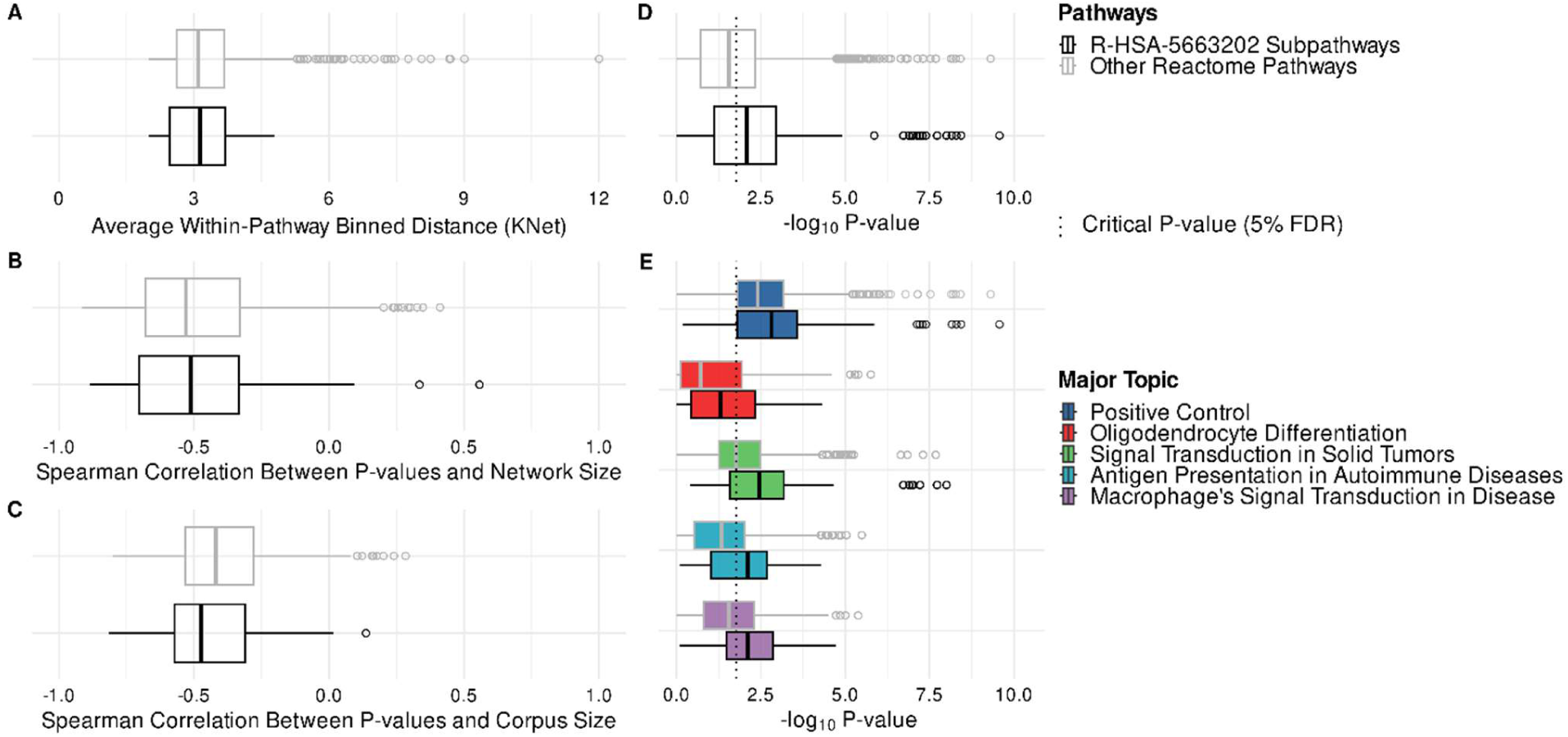
Constitutively enriched subpathways of *Diseases of signal transduction by growth factor receptors and second messengers* (R-HSA-5663202). **A**: differences in network distances between genes belonging to R-HSA-5663202 subpathways and other Reactome pathways, based on STRING’s reference physical network (FDR-adjusted p-value = 0.27, Mann-Whitney U test). The binned network distances are used by KNet to compute a topology-based pathway enrichment. **B**: differences in Spearman correlations between KNet p-values and network size, in R-HSA-5663202 subpathways and other Reactome pathways ( FDR-adjusted p-value = 0.79 , Mann-Whitney U test). **C**: differences in Spearman correlations between KNet p-values and corpus size, in R-HSA-5663202 subpathways and other Reactome pathways (FDR-adjusted p-value = 0.23, Mann-Whitney U test). **D**: differences in p-value distributions between R-HSA-5663202 subpathways and other pathways, across all case studies ( FDR-adjusted p-value = 6.5 · 10^–5^ , mixed model ANOVA). **E**: differences in p-value distributions between R-HSA-5663202 subpathways and other pathways, for each major topic ( FDR-adjusted p-value (Positive Control) = 0.04 – Mann-Whitney U test, FDR-adjusted p-value (Oligodendrocyte Differentiation) = 1.3 · 10^–2^ , FDR-adjusted p-value (Signal Transduction in Solid Tumors) = 1.4 · 10^–4^ , FDR-adjusted p-value (Antigen Presentation in Autoimmune Diseases) = 2.3 · 10^–5^ , FDR-adjusted p-value (Macrophage’s Signal Transduction in Disease) = 3.9 · 10^–4^ – mixed model ANOVA). See **Supp. Information** for details on te test statistics.

